# Effects of focal cortical cooling on somatosensory evoked potentials in rats

**DOI:** 10.1101/2023.12.28.573535

**Authors:** Mizuho Gotoh, Shinnosuke Dezawa, Ichiro Takashima, Shinya Yamamoto

**Affiliations:** Department of Information Technology and Human Factors, National Institute of Advanced Industrial Science and Technology (AIST), Tsukuba, Japan; Graduate School of Comprehensive Human Sciences, University of Tsukuba, Tsukuba Japan; Department of Rehabilitation for Brain Functions, Research Institute of National Rehabilitation Center for Persons with Disabilities, Tokorozawa, Japan; Faculty of Medical and Health Sciences, Tsukuba International University, Tsuchiura, Japan; Department of Informatics and Electronics, Daiichi Institute of Technology, Tokyo, Japan

**Keywords:** somatosensory evoked potential, temperature, focal brain cooling, gamma-aminobutyric acid, electrophysiology, rat

## Abstract

Although the focal brain cooling technique is widely used to examine brain function, the effects of cortical temperature at various levels on sensory information processing and neural mechanisms remain underexplored. To elucidate the mechanisms of temperature modulation in somatosensory processing, this study aimed to examine how P1 and N1 deflections of somatosensory evoked potentials (SEPs) depend on cortical temperature and how excitatory and inhibitory inputs contribute to this temperature dependency. SEPs were generated through electrical stimulation of the contralateral forepaw in anesthetized rats. The SEPs were recorded while cortical temperatures were altered between 17–38 °C either without any antagonists, with a gamma-aminobutyric acid type A (GABA_A_) receptor antagonist (gabazine), with aminomethylphosphonic acid (AMPA) receptor antagonist (NBQX), or with *N*-Methyl-D-aspartic acid (NMDA) receptor antagonist ([R]-CPP). The effects of different gabazine concentrations (0, 1, and 10 µM) were examined in the 35–38 °C range. The P1/N1 amplitudes and their peak-to-peak differences plotted against cortical temperature showed an inverted U relationship with a maximum at approximately 27.5 °C when no antagonists were administered. The negative correlation between these amplitudes and temperatures of ≥27.5 °C plateaued after gabazine administration, which occurred progressively as the gabazine concentration increased. In contrast, the correlation remained negative after the administration of NBQX and (R)-CPP. These results suggest that GABAergic inhibitory inputs contribute to the negative correlation between SEP amplitude and cortical temperature around the physiological cortical temperature.

**Highlights:**

- Focal cortical cooling altered somatosensory evoked potentials (SEPs).
- SEP amplitude was negatively correlated with cortical temperatures of 27.5−38.0 °C.
- GABA_A_R but not AMPAR nor NMDAR antagonists eliminated the negative correlation.
- GABAergic signaling is involved in the temperature dependency of SEPs.

## 1. Introduction^1^

The focal brain cooling (FBC) technique has been widely used in neuroscience to inactivate various brain areas (Bauer and Fuster, 1976; Fuster and Bauer, 1974; Fuster et al., 1981; Hupé et al., 1998; Lomber and Malhotra, 2008; Moseley et al., 1972; Payne et al., 1996; Takei et al., 2021; Trendelenburg, 1911). The FBC technique was first developed more than 100 years ago, whereby motor function was inactivated by cooling the motor cortex (Trendelenburg, 1911).

Since then, this technique has contributed significantly to the exploration of various neural mechanisms. This technique has also been applied to treat epilepsy in humans (Karkar et al., 2002; Nomura et al., 2017; Sartorius and Berger, 1998). Although previous neuroscience research has developed and utilized a variety of other inactivation techniques, including the physical removal (Deuel and Mishkin, 1977; Johnson et al., 2015; Minamimoto et al., 2010), pharmacological manipulation (Hikosaka et al., 1985; Hikosaka and Wurtz, 1985a; Hikosaka and Wurtz, 1985b; Newsome et al., 1985; Schwarcz et al., 1979), light-induced lesion (Nakata et al., 2018; Nakata et al., 2021; Suzuki et al., 2012), and optogenetic techniques (Chow et al., 2010; Han and Boyden, 2007; Han et al., 2011), the FBC technique has an advantage in that it can inactivate wide brain areas simultaneously in a reversible manner.

Somatosensory evoked potentials (SEPs) are field potentials generated in the somatosensory cortex in response to stimulation of the skin or peripheral nerves (e.g., the median and radial nerves). SEPs are useful in examining neural circuits and their plasticity (All et al., 2021; Dezawa et al., 2021; Franceschini et al., 2008). In addition, SEPs are used clinically to evaluate the somatosensory pathway in patients with neurological disorders and monitor the condition of patients during surgery (Carter and Butt, 2005; Cruccu et al., 2008; Florence et al., 2004; MacDonald et al., 2019). The SEPs are composed of alternating positive and negative deflections. The SEPs recorded from the primary somatosensory cortex in rats include two components: the first positive deflection after the stimulation is termed as P1, and the subsequent negative deflection is termed as N1 (Bruyns-Haylett et al., 2017; Di and Barth, 1991; Franceschini et al., 2008; Freeman and Sohmer, 1996; Jellema et al., 2004). Each deflection corresponds to a different somatosensory process in each phase.

The FBC technique has been used to examine sensory information processing. For example, unilateral cooling of the superior colliculus or the temporo-occipito-parietal junction induces visual hemineglect in cats (Payne et al., 1996). Bilateral cooling of the posterior part of the non-primary auditory cortex impairs sound localization, whereas cooling its anterior part impairs sound identification in cats (Lomber and Malhotra, 2008). In these studies, the temperature of the sensory cortices was sufficiently decreased (≤20 °C). Regarding somatosensory processing, previous studies have measured SEPs while cooling the somatosensory cortex. For example, Schwerdtfeger et al. (Schwerdtfeger et al., 1999) showed that the peak-to-peak amplitudes between P1 and N1 increased while cooling the somatosensory cortex in rats. Bindman et al. (Bindman et al., 1963) also reported that cooling the cortical surface over the somatosensory cortex increased N1 amplitudes in rats. Although these findings seemingly contradict the assumption that cooling the cortex attenuates SEPs, the estimated decrease in cortical temperature in these SEP studies was smaller than that in the FBC studies (approximately 10 °C from the physiological temperature at a depth of approximately 1 mm). Hence, the extent to which broader fluctuations in cortical temperature influence each deflection of SEP amplitudes remains unknown.

A recent study proposed that the net effects of brain temperature on neural activity involve a trade-off between excitatory and inhibitory inputs. Gotoh et al. (Gotoh et al., 2020) recorded evoked potentials in the frontal cortex that were triggered by stimulation of the mesocortical pathway. Their results showed that the increase in amplitude by focal cortical cooling was caused by a decrease in the contribution of inhibitory inputs compared to that of excitatory inputs. However, the contributions of excitatory and inhibitory inputs to the temperature dependence of sensory information processing are underexplored. In particular, elucidating their effects on each component of the SEPs (i.e., P1 and N1) would contribute significantly to a better understanding of the role of temperature modulation in sensory information processing.

To address these issues, the present study examined: (1) how the amplitudes of P1 and N1 deflections of SEPs (as well as their peak-to-peak voltages) depend on the temperature of the somatosensory cortex, and (2) how their temperature dependencies are affected by glutamate (aminomethylphosphonic acid [AMPA] and *N*-Methyl-D-aspartic acid [NMDA]) and gamma-aminobutyric acid (GABA) receptor antagonists.

## 2. Results

### 2.1 Experiment 1: Effects of cortical temperature on SEPs

We recorded SEPs in the primary somatosensory cortex that were elicited by electrical stimulation of the contralateral forepaw while altering the local cortical temperature (Fig. 1; *n* = 24 rats). The average waveforms of the SEPs from 24 rats at various temperatures are shown in Fig. 2A. The SEP waveforms changed depending on cortical temperature. To evaluate the effects of cortical temperature on the SEPs, we focused on the initial positive deflection (P1) and subsequent negative deflection (N1) (Fig. 2B, see also *4.8 Data analysis*).

**Figure 1.**
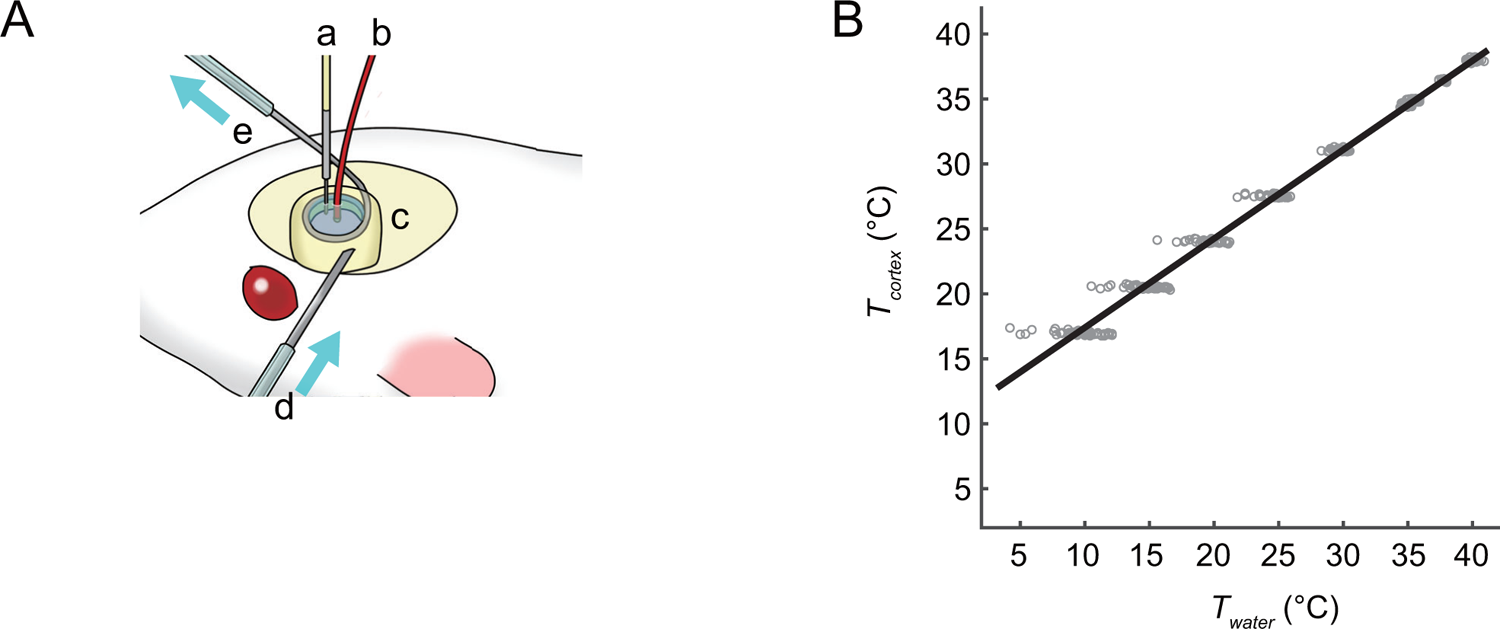
Experimental procedures. **(A)** Schematic illustration of the experimental setup. The right forepaw was stimulated, and somatosensory evoked potentials (SEPs) were recorded from the contralateral somatosensory cortex in anesthetized rats. The thermal control chamber was placed above the somatosensory cortex. a: thermocouple electrode, b: recording ball electrode, c: chamber; d: influx of circulating (temperature-controlled) water, e: efflux of circulating water. **(B)** Regulation of the cortical temperature by the thermal control chamber. Cortical temperatures (*T_cortex_*), measured at 1 mm from the surface during experiments without the antagonist, are plotted against the temperature of the circulating water (*T_water_*) through a stainless-steel tube inside the chamber and fitted by a linear function (*T_cortex_* = 0.068 × *T_water_* + 10.56, *R*^2^ = 0.99, *n* = 32 animals).

**Figure 2.**
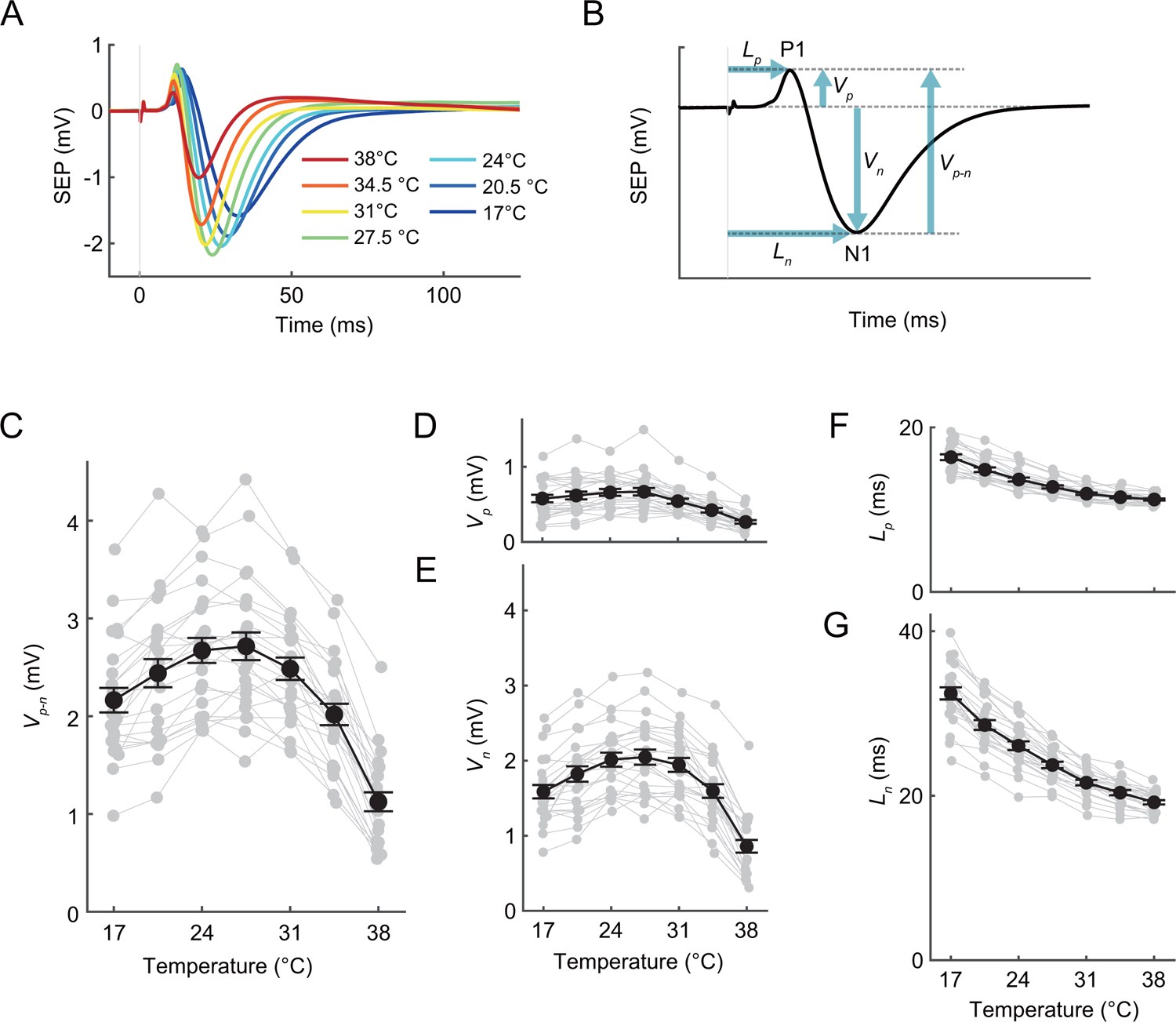
Effects of local cortical temperature on somatosensory evoked potentials (SEPs) (Experiment 1). **(A)** Averaged SEP waveforms at various temperatures from 24 animals. The time 0 indicates the onset of the electrical stimulation. **(B)** Indices of SEPs. At each temperature in each animal, the SEPs were recorded in 30 trials. After the exclusion of data from trials in which spontaneous activity was observed close to or during stimulation, we averaged the SEP data at each temperature in each animal to determine the SEP indices. The peak amplitudes of the positive peak (*V_p_*) and negative peak (*V_n_*) were defined as the absolute voltage of the respective peaks. The peak-to-peak amplitude (*V_p-n_*) was defined as the difference between positive and negative peaks. The peak latencies of the positive peak (*L_p_*) and negative peak (*L_n_*) were defined as the time between the onset of stimulation (0 ms) and the respective peaks. **(C– E)** Effects of cortical temperature on *V_p-n_* (C), *V_p_* (D), and *V_n_* (E). Amplitudes are plotted against the cortical temperatures (average data: filled circles and thick lines, individual data: semitransparent circles and thin lines; *n* = 24 animals). Error bars indicate mean ± standard error. **(F–G)** Effects of cortical temperature on *L_p_*(F) and *L_n_* (G). Peak latencies are plotted against cortical temperatures (average data: filled circles and thick lines, individual data: semitransparent circles and lines; *n* = 24 animals). Error bars indicate mean ± SE.

The relationship between temperature and *V_p-n_* exhibited a non-monotonic inverted-U shape (Fig. 2C). The *V_p-n_* was negatively correlated with the cortical temperature at ≥27.5 °C (*R* = – 0.70, *P* < 10^−5^; Supplementary Table 1), while it was positively correlated at ≤27.5 °C (*R* = 0.31, *P* = 0.0019; Supplementary Table 1). A two-way ANOVA (temperature by animal) revealed significant main effects for temperature (Table 1).

**Table 1.** Results of two-way ANOVAs (temperature by animal) for amplitudes (*V_p-n_, V_p_*, and *V_n_*) in Experiment 1.

| Effect | Sum Sq. | *d. f.* | Mean Sq. | *F* | *p* |
| --- | --- | --- | --- | --- | --- |
| Temperature | 43.49 | 6 | 7.25 | 84.2 | $1.2\times10^{-43}$ |
| Animal | 47.37 | 23 | 2.06 | 23.9 | $2.8\times10^{-37}$ |

$$V_p$$
| Effect | Sum Sq. | *d. f.* | Mean Sq. | *F* | *p* |
| --- | --- | --- | --- | --- | --- |
| Temperature | 3.05 | 6 | 0.51 | 40.9 | $2.4\times10^{-28}$ |
| Animal | 5.12 | 23 | 0.22 | 17.9 | $8.0\times10^{-31}$ |

$$V_n$$
| Effect | Sum Sq. | *d. f.* | Mean Sq. | *F* | *p* |
| --- | --- | --- | --- | --- | --- |
| Temperature | 24.56 | 6 | 4.09 | 83.0 | $2.5\times10^{-43}$ |
| Animal | 27.04 | 23 | 1.18 | 23.8 | $3.3\times10^{-37}$ |

The *V_p_* was negatively correlated with the cortical temperatures at ≥27.5 °C (*R* = –0.65, *P* < 10^−5^; Supplementary Table 1), but was not significantly correlated with the cortical temperatures at ≤27.5 °C (*R* = 0.15, *P* = 0.15; Supplementary Table 1, Fig. 2D). The *V_n_* was negatively correlated with the cortical temperatures at ≥27.5 °C (*R* = –0.67, *P* < 10^−5^; Supplementary Table 1) and was positively correlated with the cortical temperatures at ≤27.5 °C (*R* = 0.36, *P* = 0.00034; Supplementary Table 1, Fig. 2E). Two-way ANOVAs (temperature by animal) for positive and negative peaks revealed significant main effects for temperature (Table 1).

The latencies of the SEPs also changed depending on cortical temperature. Peak latency for P1 (*L_p_*) and N1 (*L_n_*) was defined as the time between the onset of stimulation and the positive and negative peaks, respectively (Fig. 2B, see *4.8 Data analysis*). Both *L_p_*and *L_n_* decreased monotonically as cortical temperature increased (Fig. 2F–G). A two-way ANOVA (temperature by animal) for negative and positive peaks revealed significant main effects for temperature (Table 2).

**Table 2.** Results of two-way ANOVAs (temperature by animal) for latencies (*L_p_* and *L_n_*) in Experiment 1.

| $L_p$ | | | | | |
| --- | --- | --- | --- | --- | --- |
| Effect | Sum Sq. | *d. f.* | Mean Sq. | *F* | *p* |
| Temperature | 516.53 | 6 | 86.09 | 229.6 | $3.2\times 10^{-69}$ |
| Animal | 139.72 | 23 | 6.07 | 16.2 | $9.3\times 10^{-29}$ |

| $L_n$ | | | | | |
| --- | --- | --- | --- | --- | --- |
| Effect | Sum Sq. | *d. f.* | Mean Sq. | *F* | *p* |
| Temperature | 3277.36 | 6 | 546.23 | 343.2 | $2.5\times 10^{-80}$ |
| Animal | 667.00 | 23 | 29.00 | 18.2 | $3.1\times 10^{-31}$ |

### 2.2 Experiment 2: Effects of gamma-aminobutyric acid type A (GABA_A_) receptor antagonist

To examine how the contribution of GABAergic inputs to the temperature dependency of SEPs, we administered the GABA_A_ receptor antagonist gabazine (10 µM, 100 µL) to the chamber in 8 of 24 rats after Experiment 1 (Fig. 3A). Gabazine administration increased *V_p-n_*. In particular, an increase in *V_p-n_* at higher temperatures resulted in the disappearance of the negative correlation at ≥27.5 °C observed in Experiment 1 (Fig. 3B; *R* = –0.11, *P* = 0.54; Supplementary Table 2). A three-way ANOVA for *V_p-n_* (antagonist by temperature and by animal) revealed significant main effects for the antagonist and temperature and significant interaction between the antagonist and temperature (Table 3). These results suggest that GABAergic inhibitory inputs contribute to the negative correlation between SEP amplitude and cortical temperature around the physiological cortical temperature.

**Figure 3.**
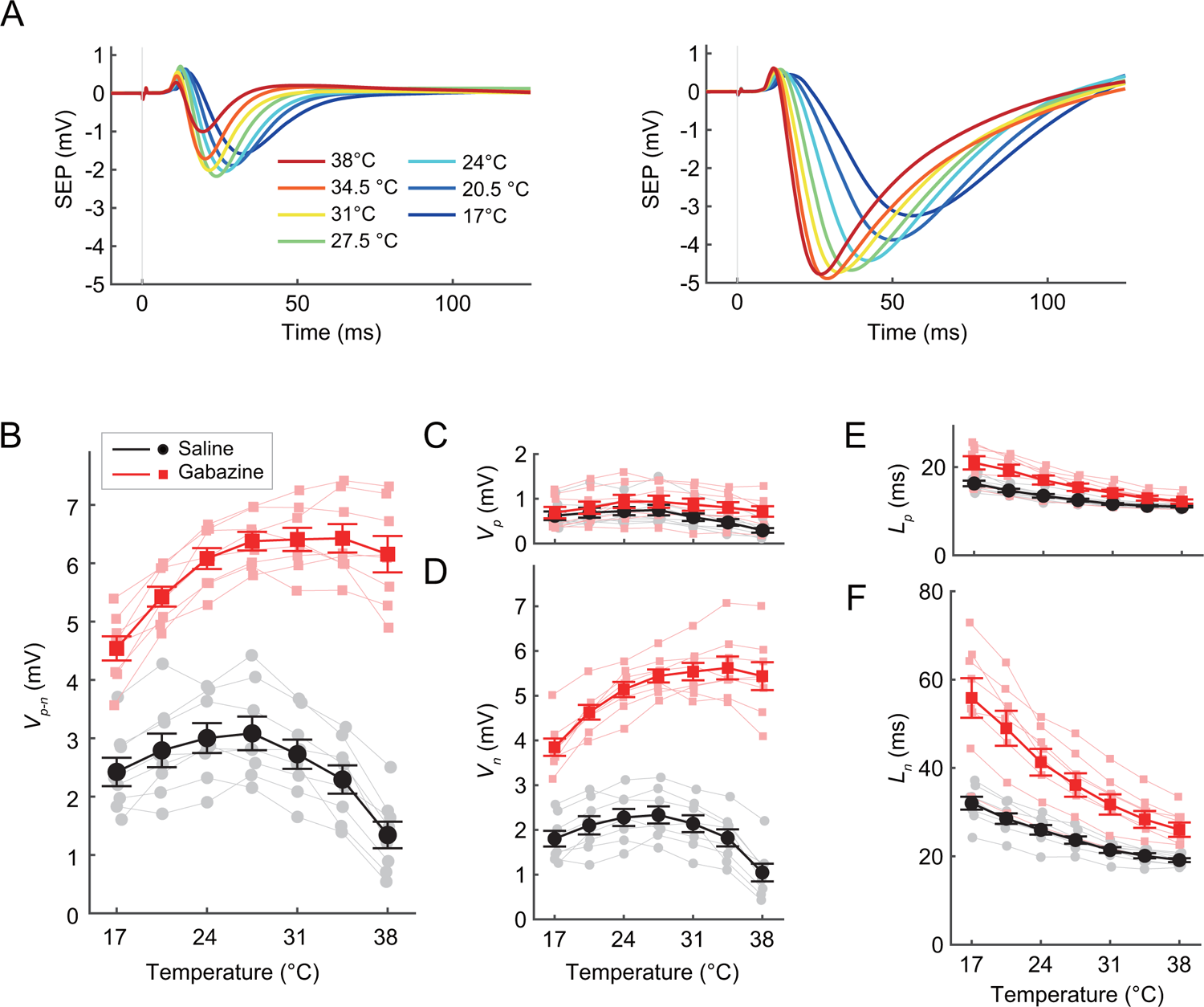
Effects of the gamma-aminobutyric acid type A (GABA_A_) receptor antagonist, gabazine, on the temperature dependency of somatosensory evoked potentials (SEPs) (Experiment 2). **(A)** Averaged SEP waveforms before (left panel) and after 10 µM gabazine administration (right panel) at various temperatures from eight animals. **(B–F)** Effects of cortical temperature on *V_p-n_* (B), *V_p_* (C), *V_n_*(D), *L_p_* (E), and *L_n_* (F) before (black circles and lines) and after (red squares and lines) gabazine administration (*n* = 8 animals). Error bars indicate mean ± standard error.

**Table 3.** Results of three-way ANOVAs (antagonist by temperature and by animal) for amplitudes (*V_p-n_, V_p_*, and *V_n_*) in Experiment 2.

| Effect | Sum Sq. | *d. f.* | Mean Sq. | *F* | *p* |
| --- | --- | --- | --- | --- | --- |
| Antagonist | 322.38 | 1 | 322.38 | 3502.1 | $4.4 \times 10^{-42}$ |
| Temperature | 20.55 | 6 | 3.42 | 37.2 | $2.8 \times 10^{-15}$ |
| Animal | 21.20 | 7 | 3.03 | 32.9 | $4.5 \times 10^{-15}$ |
| Antagonist×Temperature | 19.85 | 6 | 3.31 | 35.9 | $5.2 \times 10^{-15}$ |
| Antagonist×Animal | 12.90 | 7 | 1.84 | 20.0 | $1.7 \times 10^{-11}$ |
| Temperature×Animal | 6.22 | 42 | 0.15 | 1.6 | 0.064 |

$$V_p$$
| Effect | Sum Sq. | *d. f.* | Mean Sq. | *F* | *p* |
| --- | --- | --- | --- | --- | --- |
| Antagonist | 1.50 | 1 | 1.50 | 166.8 | $3.2 \times 10^{-16}$ |
| Temperature | 1.34 | 6 | 0.22 | 25.0 | $2.2 \times 10^{-12}$ |
| Animal | 7.46 | 7 | 1.07 | 118.8 | $1.4 \times 10^{-25}$ |
| Antagonist×Temperature | 0.37 | 6 | 0.06 | 6.9 | $3.8 \times 10^{-5}$ |
| Antagonist×Animal | 1.35 | 7 | 0.19 | 21.5 | $5.3 \times 10^{-12}$ |
| Temperature×Animal | 0.89 | 42 | 0.02 | 2.4 | $3.2 \times 10^{-3}$ |

$$V_n$$
| Effect | Sum Sq. | *d. f.* | Mean Sq. | *F* | *p* |
| --- | --- | --- | --- | --- | --- |
| Antagonist | 279.95 | 1 | 279.95 | 4155.4 | $1.2 \times 10^{-43}$ |
| Temperature | 14.33 | 6 | 2.39 | 35.5 | $6.5 \times 10^{-15}$ |
| Animal | 13.68 | 7 | 1.95 | 29.0 | $3.9 \times 10^{-14}$ |
| Antagonist×Temperature | 14.85 | 6 | 2.48 | 36.8 | $3.5 \times 10^{-15}$ |
| Antagonist×Animal | 11.81 | 7 | 1.69 | 25.0 | $4.6 \times 10^{-13}$ |
| Temperature×Animal | 3.75 | 42 | 0.09 | 1.3 | 0.18 |

Gabazine administration increased individual peak amplitudes (*V_p_*and *V_n_*). The correlations between peak amplitudes (*V_p_*and *V_n_*) and temperature at ≥27.5 °C were not significantly different from 0 in Experiment 2 (Fig. 3C–D, *V_p_*: *R* = –0.23, *P* = 0.21, *V_n_*: *R* = 0.010, *P* = 0.96; Supplementary Table 2). A three-way ANOVA for *V_p_* and *V_n_* (antagonist by temperature and by animal) revealed significant main effects for the antagonist and temperature and significant interaction between the antagonist and temperature (Table 3).

To evaluate the contribution of GABAergic inputs, we calculated the difference and ratio between the amplitudes before and after gabazine administration at each cortical temperature level. Both the difference and the ratio of the amplitudes increased at higher temperatures for *V_p-n_, V_p_*, and *V_n_* (Fig. S1). The correlation between *difference*(*V_p-n_*) and temperature, that between *difference*(*V_p_*) and temperature, and that between *difference*(*V_n_*) and temperature were significantly positive (Supplementary Table 3). The correlation between *ratio*(*V_p-n_*) and temperature, that between *ratio*(*V_p_*) and temperature, and that between *ratio*(*V_n_*) and temperature were also significantly positive (Supplementary Table 4). These results suggest that the effect of gabazine (i.e., an increase in *V_p-n_, V_p_*, and *V_n_*) was greater at higher temperatures.

Gabazine administration influenced the latency. In Experiment 2, both *L_p_* and *L_n_* exhibited values higher than those recorded in Experiment 1; nonetheless, they decreased monotonically as the temperature rose (Fig. 3E–F). A three-way ANOVA (antagonist by temperature and by animal) revealed significant main effects for the antagonist and temperature and significant interaction between the antagonist and temperature (Table 4). These results suggest that GABAergic inputs increase the peak latencies of SEPs.

**Table 4.** Results of three-way ANOVAs (antagonist by temperature and by animal) for latencies (*L_p_*and *L_n_*) in Experiment 2.

| Effect | Sum Sq. | *d. f.* | Mean Sq. | *F* | *p* |
| --- | --- | --- | --- | --- | --- |
| Antagonist | 254.27 | 1 | 254.27 | 466.2 | $2.3 \times 10^{-24}$ |
| Temperature | 652.05 | 6 | 108.68 | 199.3 | $3.3 \times 10^{-29}$ |
| Animal | 252.77 | 7 | 36.11 | 66.2 | $1.2 \times 10^{-20}$ |
| Antagonist×Temperature | 43.50 | 6 | 7.25 | 13.3 | $2.2 \times 10^{-08}$ |
| Antagonist×Animal | 74.50 | 7 | 10.64 | 19.5 | $2.5 \times 10^{-11}$ |
| Temperature×Animal | 72.23 | 42 | 1.72 | 3.2 | $1.6 \times 10^{-4}$ |

$L_n$
| Effect | Sum Sq. | *d. f.* | Mean Sq. | *F* | *p* |
| --- | --- | --- | --- | --- | --- |
| Antagonist | 5433.87 | 1 | 5433.87 | 1159.4 | $3.2 \times 10^{-32}$ |
| Temperature | 5940.22 | 6 | 990.04 | 211.2 | $1.0 \times 10^{-29}$ |
| Animal | 2361.26 | 7 | 337.32 | 72.0 | $2.5 \times 10^{-21}$ |
| Antagonist×Temperature | 958.68 | 6 | 159.78 | 34.1 | $1.3 \times 10^{-14}$ |
| Antagonist×Animal | 881.81 | 7 | 125.97 | 26.9 | $1.4 \times 10^{-13}$ |
| Temperature×Animal | 480.69 | 42 | 11.45 | 2.4 | $2.3 \times 10^{-3}$ |

### 2.3 Experiment 3: Effects of different concentrations of GABA_A_ receptor antagonist around the physiological temperature

To evaluate the effects of concentration of gabazine around the physiological temperature, we recorded the SEPs at 35 °C, 36.5 °C, and 38 °C with the chamber filled with saline, 1 µM gabazine, and 10 µM gabazine (Fig. 4). As the gabazine concentration increased, *V_p-n_, V_p_*, and *V_n_*increased, and the negative correlation between the amplitude and temperature plateaued (Fig. 4A–C). Three-way ANOVAs for *V_p-n_, V_p_*, and *V_n_* (antagonist by temperature and by animal) revealed significant main effects for the antagonist and temperature and significant interaction between the antagonist and temperature (Table 5).

**Figure 4.**
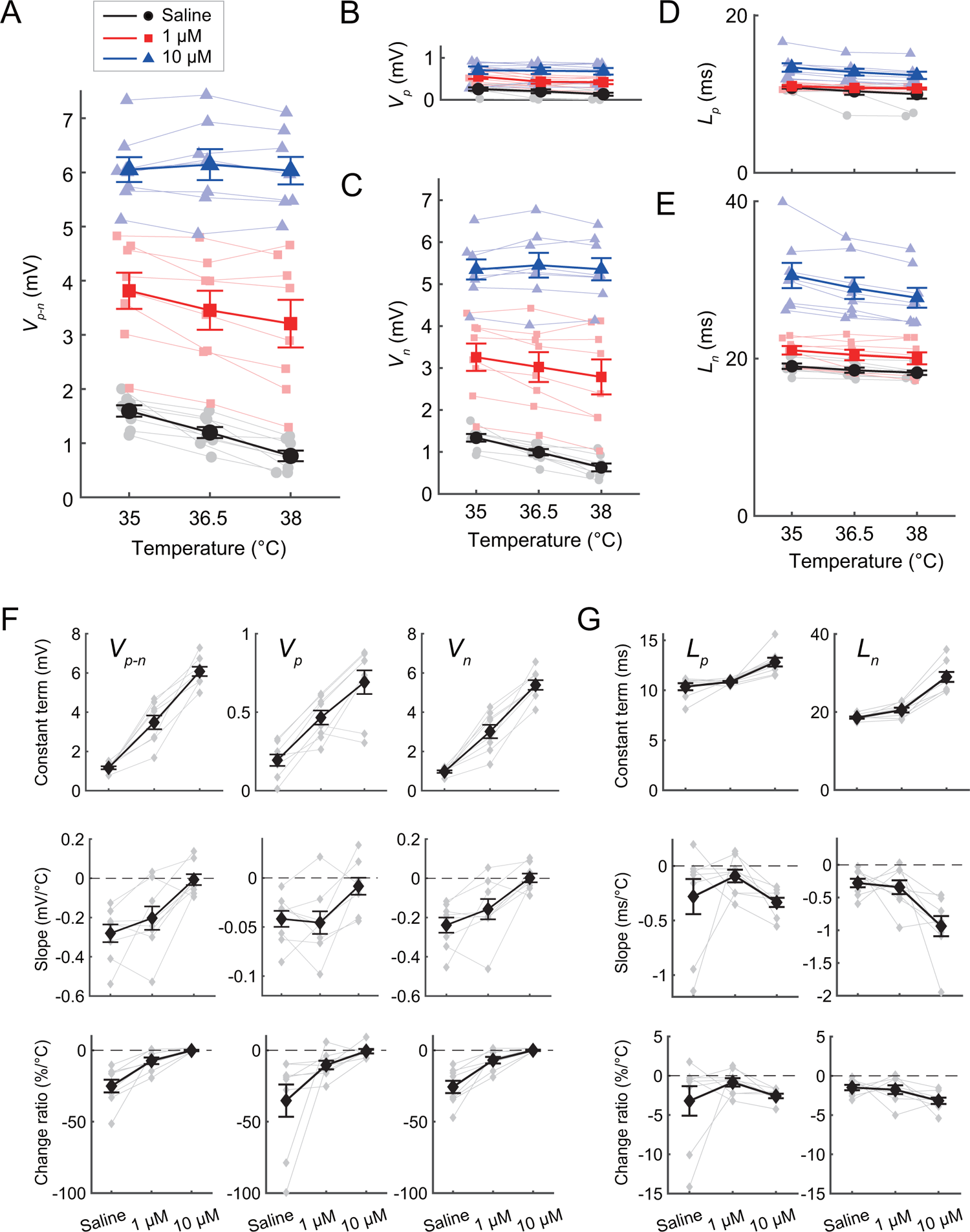
Effects of the concentration of the gamma-aminobutyric acid type A (GABA_A_) receptor antagonist, gabazine, on the temperature dependency of somatosensory evoked potentials (SEPs) (Experiment 3). **(A–E)** Effects of cortical temperature on *V_p-n_* (A), *V_p_* (B), *V_n_*(C), *L_p_* (D), and *L_n_* (E) in the saline (black circles and lines), 1-µM gabazine (red squares and lines), and 10-µM gabazine (blue triangles and lines) condition (*n* = 8 animals). Error bars indicate mean ± standard error. **(F)** Linear regression of the amplitudes. The constant terms (top panels; the estimated amplitudes at 36.5 °C), slopes (middle panels), and the change ratios (bottom panels) of the fitted linear models for *V_p-n_, V_p_*, and *V_n_* are plotted against the gabazine concentration (black diamonds and lines; *n* = 8 animals). **(G)** Linear regression of the latencies. The constant terms (top panels; the estimated latencies at 36.5 °C), slopes (middle panels), and the change ratios (bottom panels) of the fitted linear models for *L_p_* and *L_n_* are plotted against the gabazine concentration (black diamonds and lines; *n* = 8 animals).

**Table 5.** Results of three-way ANOVAs (antagonist by temperature and by animal) for amplitudes (*V_p-n_, V_p_*, and *V_n_*) in Experiment 3.

| Effect | Sum Sq. | *d. f.* | Mean Sq. | *F* | *p* |
| --- | --- | --- | --- | --- | --- |
| Antagonist | 287.36 | 2 | 143.68 | 3286.2 | $6.1 \times 10^{-34}$ |
| Temperature | 2.83 | 2 | 1.41 | 32.3 | $5.3 \times 10^{-8}$ |
| Animal | 15.07 | 7 | 2.15 | 49.2 | $4.5 \times 10^{-14}$ |
| Antagonist×Temperature | 1.47 | 4 | 0.37 | 8.4 | $1.4 \times 10^{-4}$ |
| Antagonist×Animal | 20.06 | 14 | 1.43 | 32.8 | $1.2 \times 10^{-13}$ |
| Temperature×Animal | 0.93 | 14 | 0.066 | 1.5 | 0.17 |

$$V_p$$
| Effect | Sum Sq. | *d. f.* | Mean Sq. | *F* | *p* |
| --- | --- | --- | --- | --- | --- |
| Antagonist | 2.94 | 2 | 1.47 | 534.9 | $4.9 \times 10^{-23}$ |
| Temperature | 0.11 | 2 | 0.056 | 20.4 | $3.5 \times 10^{-6}$ |
| Animal | 1.26 | 7 | 0.18 | 65.3 | $1.2 \times 10^{-15}$ |
| Antagonist×Temperature | 0.044 | 4 | 0.01 | 4.0 | 0.010 |
| Antagonist×Animal | 0.49 | 14 | 0.04 | 12.8 | $1.2 \times 10^{-8}$ |
| Temperature×Animal | 0.043 | 14 | 0.0031 | 1.1 | 0.39 |

$$V_n$$
| Effect | Sum Sq. | *d. f.* | Mean Sq. | *F* | *p* |
| --- | --- | --- | --- | --- | --- |
| Antagonist | 232.40 | 2 | 116.20 | 3430.9 | $3.4 \times 10^{-34}$ |
| Temperature | 1.85 | 2 | 0.92 | 27.3 | $2.7 \times 10^{-7}$ |
| Animal | 17.73 | 7 | 2.53 | 74.8 | $2.0 \times 10^{-16}$ |
| Antagonist×Temperature | 1.08 | 4 | 0.27 | 8.0 | $2.0 \times 10^{-4}$ |
| Antagonist×Animal | 16.73 | 14 | 1.19 | 35.3 | $4.7 \times 10^{-14}$ |
| Temperature×Animal | 0.66 | 14 | 0.047 | 1.4 | 0.22 |

To evaluate the effects of cortical temperatures within the physiologically normal temperature range on SEPs, we fitted *V_p-n_, V_p_*, and *V_n_* at each concentration using a linear model of the temperatures (Supplementary Table 5; see also *4.8 Data Analysis*). The constant terms (the estimated amplitudes at 36.5 °C) of the fitted linear models increased at higher concentrations (Fig. 4F, top). The two-way ANOVA for the constant terms of *V_p-n_, V_p_*, and *V_n_* (antagonist by animal) revealed significant main effects of the antagonist (Supplementary Table 6). The slopes increased from −0.28 mV/°C to −0.0065 mV/°C in *V_p-n_*, −0.042 mV/°C to −0.0084 mV/°C in *V_p_*, −0.24 mV/°C to 0.0019 mV/°C in *V_n_* as the concentration increased (Fig. 4F, middle). Two-way ANOVA of the slopes (antagonist by animal) revealed significant main effects of the antagonist (Supplementary Table 7). The slopes in the saline and 1-µM gabazine conditions were significantly smaller than 0 but not in the 10-µM gabazine condition (*t*-test; Supplementary Table 5). The change ratios (i.e., the ratio of the slope to the constant term) also increased from −25.02 %/°C to −0.16 %/°C in *V_p-n_*, −35.23 %/°C to −0.63 %/°C in *V_p_*, −25.64 %/°C to −0.022 %/°C in *V_n_*as the concentration increased (Fig. 4F, bottom). Two-way ANOVA of the change ratio (antagonist by animal) revealed significant main effects of the antagonist (Supplementary Table 8). The change ratios in the saline and 1-µM gabazine conditions were significantly smaller than 0 but not in the 10-µM gabazine condition (*t*-test; Supplementary Table 5). The results showed that a higher concentration of gabazine had a greater effect on the peak amplitudes.

The effects of gabazine on the latencies (*L_p_* and *L_n_*) were also greater at higher concentrations (Fig. 4D–E). A three-way ANOVA (antagonist by temperature and by animal) revealed significant main effects for the antagonist and temperature (Table 6). The interaction between the antagonist and temperature was significant for *L_n_* but not for *L_p_*. We also fitted *L_p_* and *L_n_* at each concentration using a linear model of the temperatures (Supplementary Table 9). The constant terms (the estimated latencies at 36.5 °C) of the fitted linear models increased at higher concentrations (Fig. 4G, top). The two-way ANOVA for the constant terms of *L_p_*and *L_n_* (antagonist by animal) revealed significant main effects of the antagonist (Supplementary Table 10). The effect of the antagonist concentration on the slope and the change ratio of *L_p_* was less clear. The two-way ANOVA for the slope of *L_p_* (antagonist by animal) and that for the change ratio of *L_p_* (antagonist by animal) revealed no significant main effects of the antagonist (Supplementary Tables 11 and 12). The two-way ANOVA for the slope of *L_n_* (antagonist by animal) revealed significant main effects of the antagonist but that for the change ratio of *L_n_* (antagonist by animal) did not (Supplementary Tables 11 and 12). The results suggest that the latency approached the asymptote around the physiological temperature.

**Table 6.**
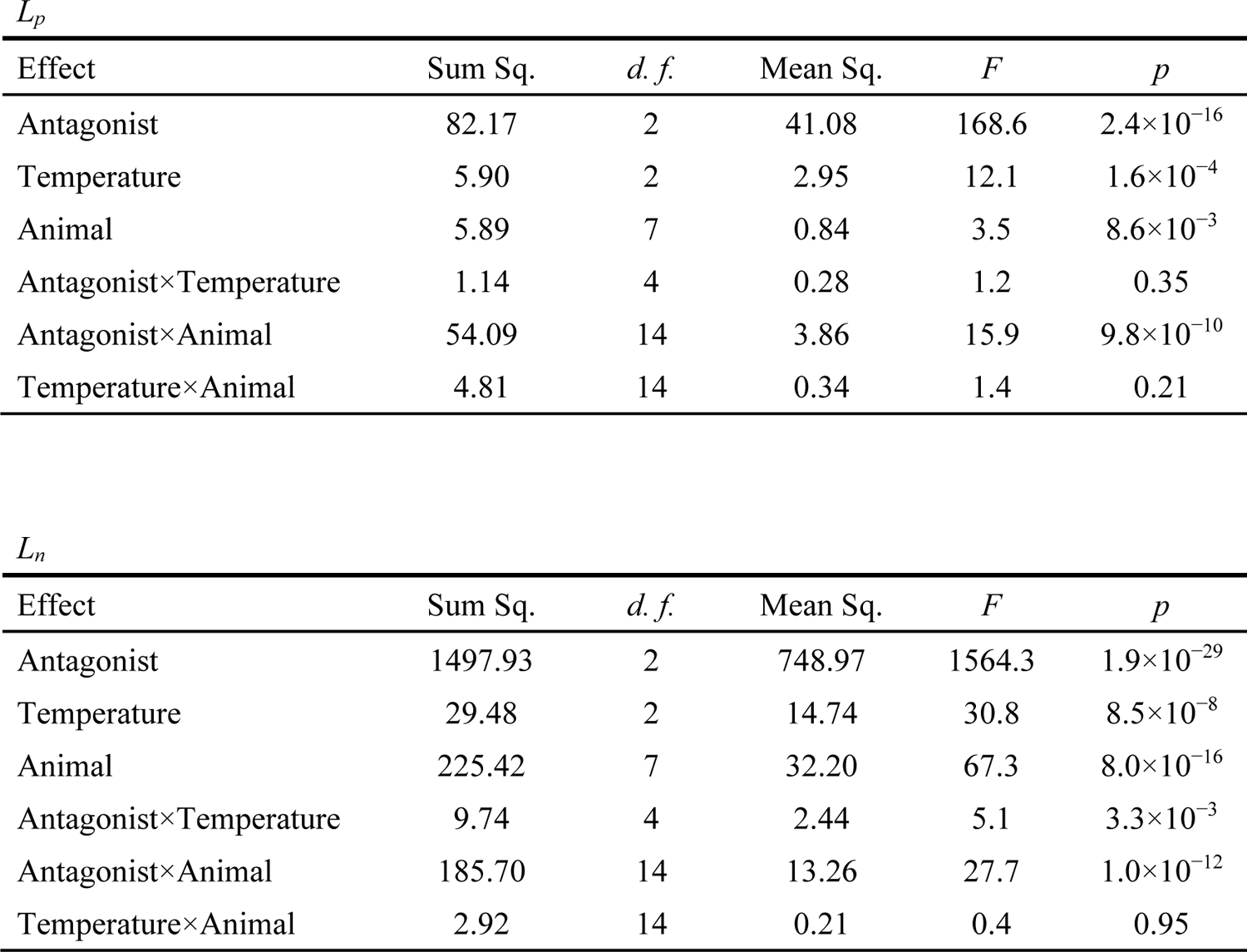
Results of three-way ANOVAs (antagonist by temperature and by animal) for latencies (*L_p_*and *L_n_*) in Experiment 3.

| Effect | Sum Sq. | *d. f.* | Mean Sq. | *F* | *p* |
| --- | --- | --- | --- | --- | --- |
| Antagonist | 82.17 | 2 | 41.08 | 168.6 | $2.4 \times 10^{-16}$ |
| Temperature | 5.90 | 2 | 2.95 | 12.1 | $1.6 \times 10^{-4}$ |
| Animal | 5.89 | 7 | 0.84 | 3.5 | $8.6 \times 10^{-3}$ |
| Antagonist×Temperature | 1.14 | 4 | 0.28 | 1.2 | 0.35 |
| Antagonist×Animal | 54.09 | 14 | 3.86 | 15.9 | $9.8 \times 10^{-10}$ |
| Temperature×Animal | 4.81 | 14 | 0.34 | 1.4 | 0.21 |

$L_n$
| Effect | Sum Sq. | *d. f.* | Mean Sq. | *F* | *p* |
| --- | --- | --- | --- | --- | --- |
| Antagonist | 1497.93 | 2 | 748.97 | 1564.3 | $1.9 \times 10^{-29}$ |
| Temperature | 29.48 | 2 | 14.74 | 30.8 | $8.5 \times 10^{-8}$ |
| Animal | 225.42 | 7 | 32.20 | 67.3 | $8.0 \times 10^{-16}$ |
| Antagonist×Temperature | 9.74 | 4 | 2.44 | 5.1 | $3.3 \times 10^{-3}$ |
| Antagonist×Animal | 185.70 | 14 | 13.26 | 27.7 | $1.0 \times 10^{-12}$ |
| Temperature×Animal | 2.92 | 14 | 0.21 | 0.4 | 0.95 |

### 2.4 Experiment 4: Effects of AMPA receptor antagonist

To examine the effects of glutamatergic inputs to AMPA receptors on SEPs, we administered the AMPA receptor antagonist NBQX (10 µM, 100 µL) to the chamber in eight of 24 rats after Experiment 1 (Fig. 5A). The administration of NBQX consistently reduced *V_p-n_* across all tested temperatures (Fig. 5B). The correlation between *V_p-n_* and temperature at ≥27.5 °C remained negative (*R* = −0.69, *P* = 0.000013; Supplementary Table 13). A three-way ANOVA (antagonist by temperature and by animal) revealed significant main effects for the antagonist and temperature. The interaction between the antagonist and temperature was not significant (Table 7). Both the difference and ratio of *V_p-n_* before and after the administration of NBQX were almost constant across temperatures (Fig. S2A). The correlation between the *difference*(*V_p-n_*) and temperature, as well as the *ratio*(*V_p-n_*) and temperature, did not significantly deviate from 0 (*difference*[*V_p-n_*]: *R* = 0.21, *P* = 0.12, *ratio*[*V_p-n_*]: *R* = 0.057, *P* = 0.68; Supplementary Tables 14– 15). This implies that the impact of NBQX (i.e., the decrease in *V_p-n_*) remained constant across temperatures.

**Figure 5.**
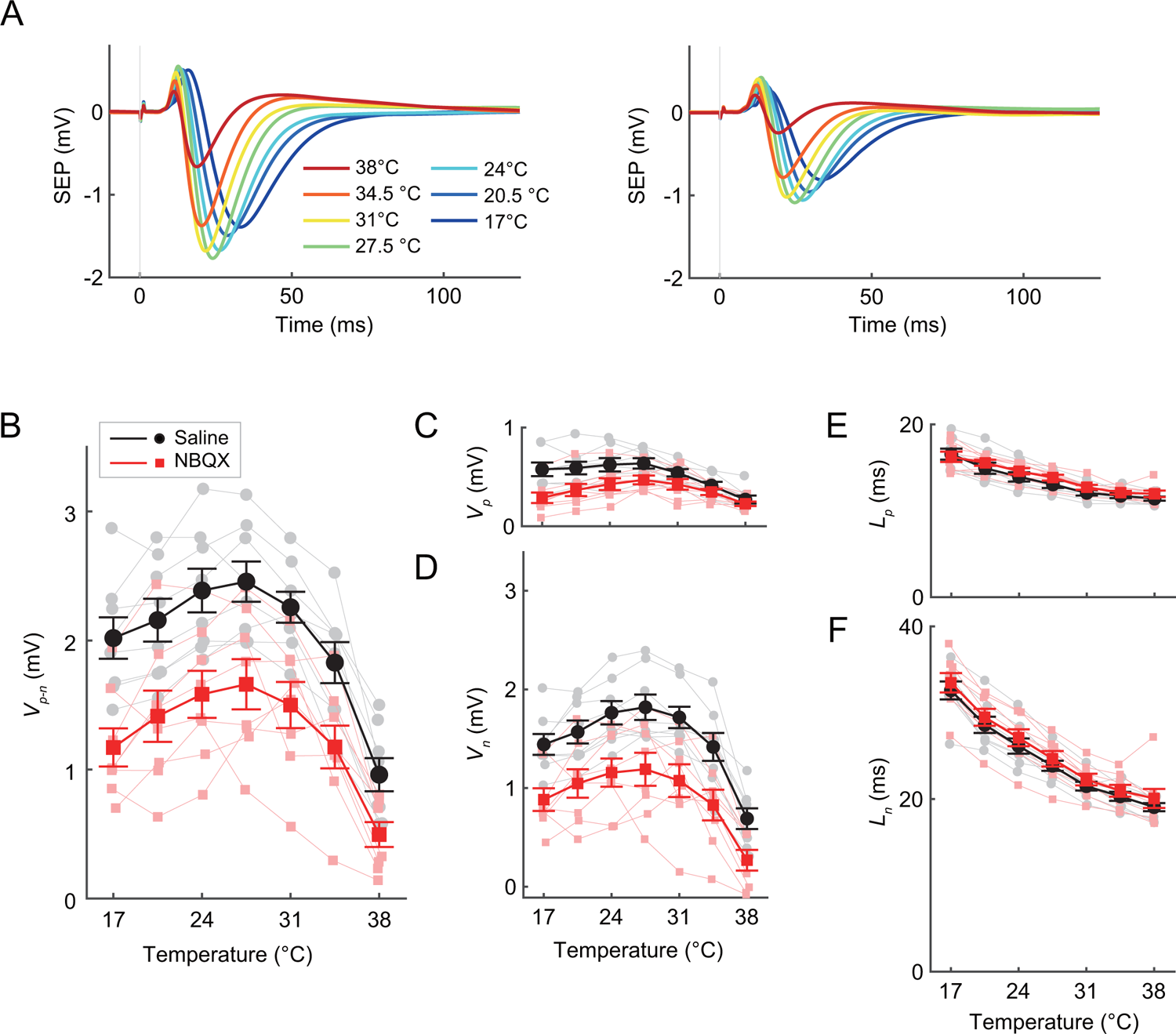
Effects of the aminomethylphosphonic acid (AMPA) receptor antagonist, NBQX, on the temperature dependency of somatosensory evoked potentials (SEPs) (Experiment 4). **(A)** Averaged SEP waveforms before (left panel) and after (right panel) NBQX administration at various temperatures from eight animals. **(B–F)** Effects of cortical temperature on *V_p-n_* (B), *V_p_*(C), *V_n_* (D), *L_p_* (E), and *L_n_*(F) before (black circles and lines) and after (red squares and lines) NBQX administration (*n* = 8 animals). Error bars indicate mean ± standard error.

**Table 7.**
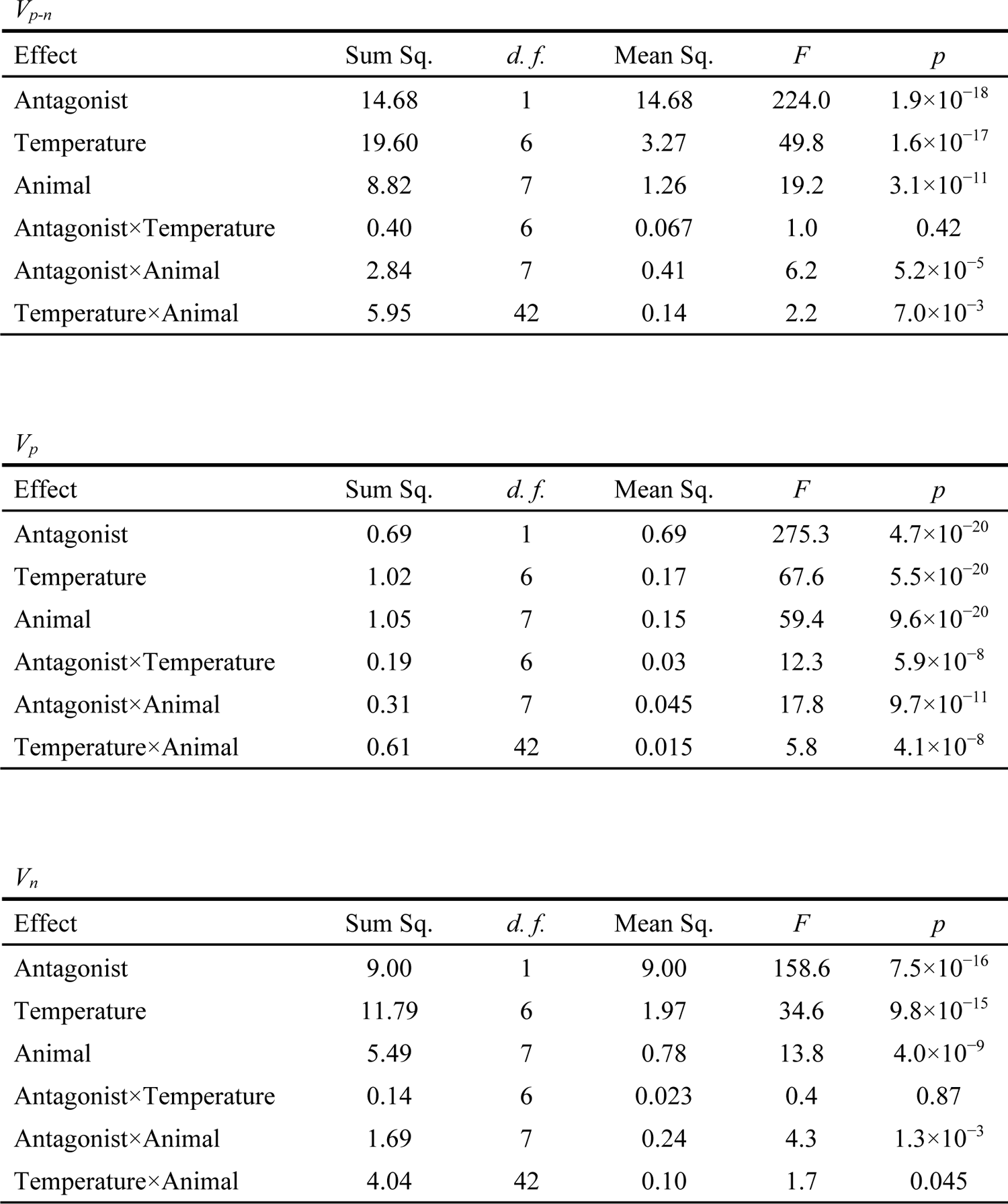
Results of three-way ANOVAs (antagonist by temperature and by animal) for amplitudes (*V_p-n_, V_p_*, and *V_n_*) in Experiment 4.

| Effect | Sum Sq. | *d. f.* | Mean Sq. | *F* | *p* |
| --- | --- | --- | --- | --- | --- |
| Antagonist | 14.68 | 1 | 14.68 | 224.0 | $1.9 \times 10^{-18}$ |
| Temperature | 19.60 | 6 | 3.27 | 49.8 | $1.6 \times 10^{-17}$ |
| Animal | 8.82 | 7 | 1.26 | 19.2 | $3.1 \times 10^{-11}$ |
| Antagonist×Temperature | 0.40 | 6 | 0.067 | 1.0 | 0.42 |
| Antagonist×Animal | 2.84 | 7 | 0.41 | 6.2 | $5.2 \times 10^{-5}$ |
| Temperature×Animal | 5.95 | 42 | 0.14 | 2.2 | $7.0 \times 10^{-3}$ |

$$V_p$$
| Effect | Sum Sq. | *d. f.* | Mean Sq. | *F* | *p* |
| --- | --- | --- | --- | --- | --- |
| Antagonist | 0.69 | 1 | 0.69 | 275.3 | $4.7 \times 10^{-20}$ |
| Temperature | 1.02 | 6 | 0.17 | 67.6 | $5.5 \times 10^{-20}$ |
| Animal | 1.05 | 7 | 0.15 | 59.4 | $9.6 \times 10^{-20}$ |
| Antagonist×Temperature | 0.19 | 6 | 0.03 | 12.3 | $5.9 \times 10^{-8}$ |
| Antagonist×Animal | 0.31 | 7 | 0.045 | 17.8 | $9.7 \times 10^{-11}$ |
| Temperature×Animal | 0.61 | 42 | 0.015 | 5.8 | $4.1 \times 10^{-8}$ |

$$V_n$$
| Effect | Sum Sq. | *d. f.* | Mean Sq. | *F* | *p* |
| --- | --- | --- | --- | --- | --- |
| Antagonist | 9.00 | 1 | 9.00 | 158.6 | $7.5 \times 10^{-16}$ |
| Temperature | 11.79 | 6 | 1.97 | 34.6 | $9.8 \times 10^{-15}$ |
| Animal | 5.49 | 7 | 0.78 | 13.8 | $4.0 \times 10^{-9}$ |
| Antagonist×Temperature | 0.14 | 6 | 0.023 | 0.4 | 0.87 |
| Antagonist×Animal | 1.69 | 7 | 0.24 | 4.3 | $1.3 \times 10^{-3}$ |
| Temperature×Animal | 4.04 | 42 | 0.10 | 1.7 | 0.045 |

Administration of NBQX decreased *V_p_* with the correlation at ≥27.5 °C remained negative (Fig. 5C; *R* = −0.63, *P* = 0.000095; Supplementary Table 13). A three-way ANOVA (antagonist by temperature and by animal) revealed significant main effects for the antagonist and temperature and the interaction between the antagonist and temperature (Table 7). The difference and ratio of *V_p_* before and after NBQX administration—referred to as *difference*(*V_p_*) and *ratio*(*V_p_*)— increased at elevated temperatures (Fig. S2B). The correlation between *difference*(*V_p_*) and temperature and that between *ratio*(*V_p_*) and temperature were significantly positive (*difference*(*V_p_*): *R* = 0.55, *P* = 0.000011, *ratio*(*V_p_*): *R* = 0.52, *P* = 0.000038; Supplementary Tables14–15), suggesting that the effect of NBQX (i.e., the decrease of *V_p_*) was larger at the lower temperatures.

Administration of NBQX also decreased *V_n_* with the correlation at ≥27.5 °C remaining negative (Fig. 5D; *R* = −0.63, *P* = 0.000097; Supplementary Table 13). A three-way ANOVA (antagonist by temperature and by animal) revealed significant main effects for the antagonist and temperature. The interaction between the antagonist and temperature was not significant (Table 7). Both the difference and ratio of *V_n_* before and after the administration of NBQX were almost constant across temperatures (Fig. S2C). The correlation between *difference*(*V_n_*) and temperature and that between *ratio*(*V_n_*) and temperature were not significantly different from 0 (*difference*(*V_n_*): *R* = 0.045, *P* = 0.74, *ratio*(*V_n_*): *R* = −0.14, *P* = 0.31; Supplementary Tables 14– 15), implying that the effect of NBQX (i.e., the decrease of *V_n_*) was constant across temperatures.

Administering NBQX caused a slight but significant increase in the peak latencies (Fig. 5E–F). *L_p_* remained negatively correlated with the cortical temperature after administering NBQX but was 0.48 ms longer on average. A three-way ANOVA (antagonist by temperature and by animal) revealed significant main effects for the antagonist, for temperature, and significant interaction between the antagonist and temperature. Administering NBQX also caused a slight but significant increase in *L_n_* by 0.85 ms on average while remaining negatively correlated with the cortical temperature. A three-way ANOVA (antagonist by temperature and by animal) revealed significant main effects for the antagonist and temperature. The interaction between the antagonist and temperature was insignificant (Table 8).

**Table 8.**
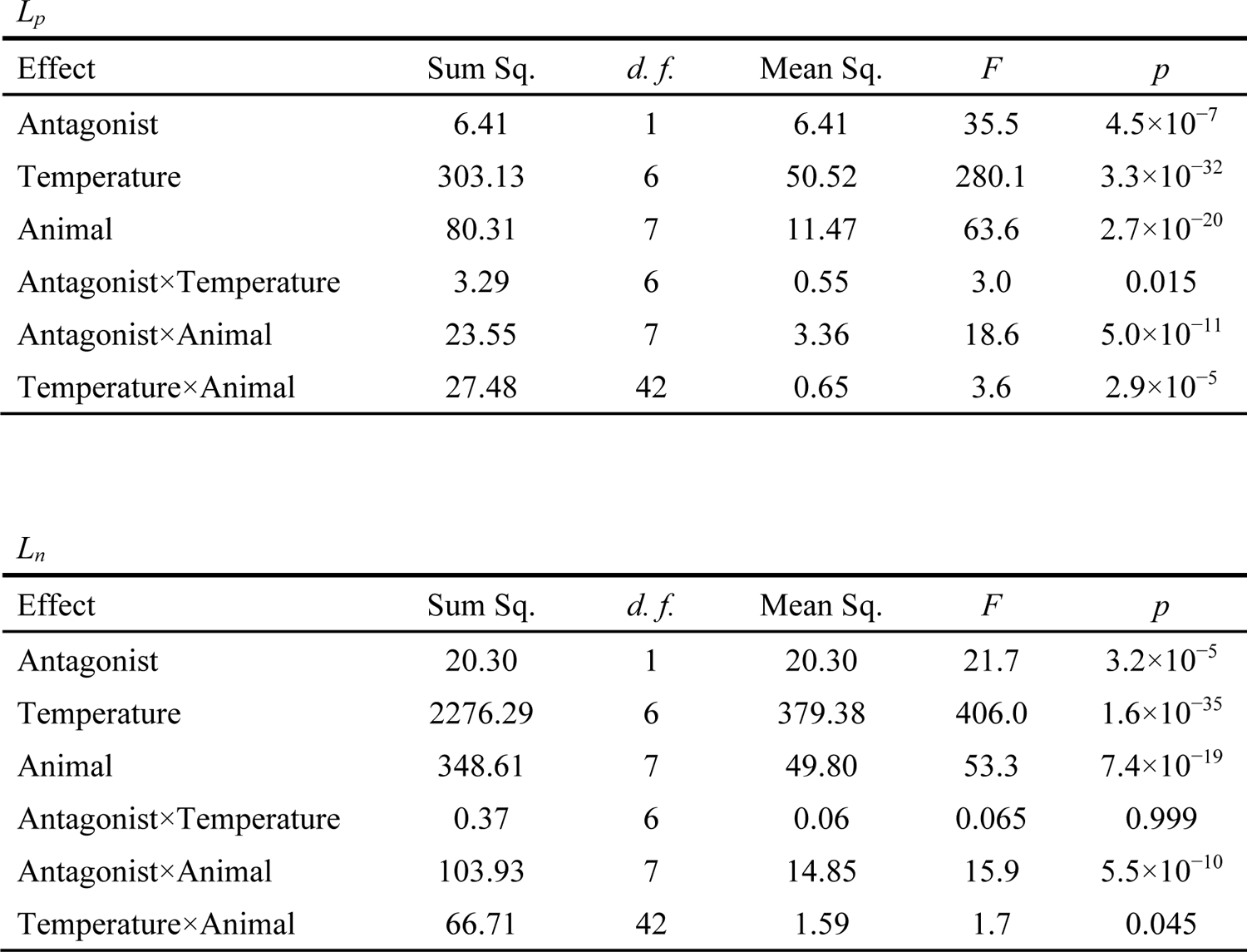
Results of three-way ANOVAs (antagonist by temperature and by animal) for latencies (*L_p_*and *L_n_*) in Experiment 4.

| Effect | Sum Sq. | *d. f.* | Mean Sq. | *F* | *p* |
| --- | --- | --- | --- | --- | --- |
| Antagonist | 6.41 | 1 | 6.41 | 35.5 | $4.5 \times 10^{-7}$ |
| Temperature | 303.13 | 6 | 50.52 | 280.1 | $3.3 \times 10^{-32}$ |
| Animal | 80.31 | 7 | 11.47 | 63.6 | $2.7 \times 10^{-20}$ |
| Antagonist×Temperature | 3.29 | 6 | 0.55 | 3.0 | 0.015 |
| Antagonist×Animal | 23.55 | 7 | 3.36 | 18.6 | $5.0 \times 10^{-11}$ |
| Temperature×Animal | 27.48 | 42 | 0.65 | 3.6 | $2.9 \times 10^{-5}$ |

$L_n$
| Effect | Sum Sq. | *d. f.* | Mean Sq. | *F* | *p* |
| --- | --- | --- | --- | --- | --- |
| Antagonist | 20.30 | 1 | 20.30 | 21.7 | $3.2 \times 10^{-5}$ |
| Temperature | 2276.29 | 6 | 379.38 | 406.0 | $1.6 \times 10^{-35}$ |
| Animal | 348.61 | 7 | 49.80 | 53.3 | $7.4 \times 10^{-19}$ |
| Antagonist×Temperature | 0.37 | 6 | 0.06 | 0.065 | 0.999 |
| Antagonist×Animal | 103.93 | 7 | 14.85 | 15.9 | $5.5 \times 10^{-10}$ |
| Temperature×Animal | 66.71 | 42 | 1.59 | 1.7 | 0.045 |

### 2.5 Experiment 5: Effects of NMDA receptor antagonist

To examine the effects of glutamatergic inputs to NMDA receptors on SEPs, we administered the NMDA receptor antagonist (R)-CPP (10 µM, 100 µL) to the chamber in eight of 24 rats after Experiment 1 (Fig. 6A). *V_p-n_* decreased after administration of (R)-CPP, particularly at higher temperatures (Fig. 6B). The correlation between *V_p-n_* and temperature at ≥27.5 °C remained negative (*R* = –0.86, *P* < 10^−5^; Supplementary Table 16). A three-way ANOVA (antagonist by temperature and by animal) revealed significant main effects for the antagonist and temperature and the interaction between the antagonist and temperature (Table 9). Both the difference and ratio of *V_p-n_* before and after the administration of (R)-CPP (that is, the *difference*(*V_p-n_*) and *ratio*(*V_p-n_*)) were smaller at higher temperatures (Fig. S3A). The correlation between *difference*(*V_p-n_*) and temperature and that between *ratio*(*V_p-n_*) and temperature were significantly negative (*difference*(*V_p-n_*): *R* = −0.52, *P* = 0.000038, *ratio*(*V_p-n_*): *R* = −0.64, *P* < 10^−5^; Supplementary Tables 17–18), suggesting that the effect of (R)-CPP (i.e., the decrease of *V_p-n_*) was larger at the higher temperatures.

**Figure 6.**
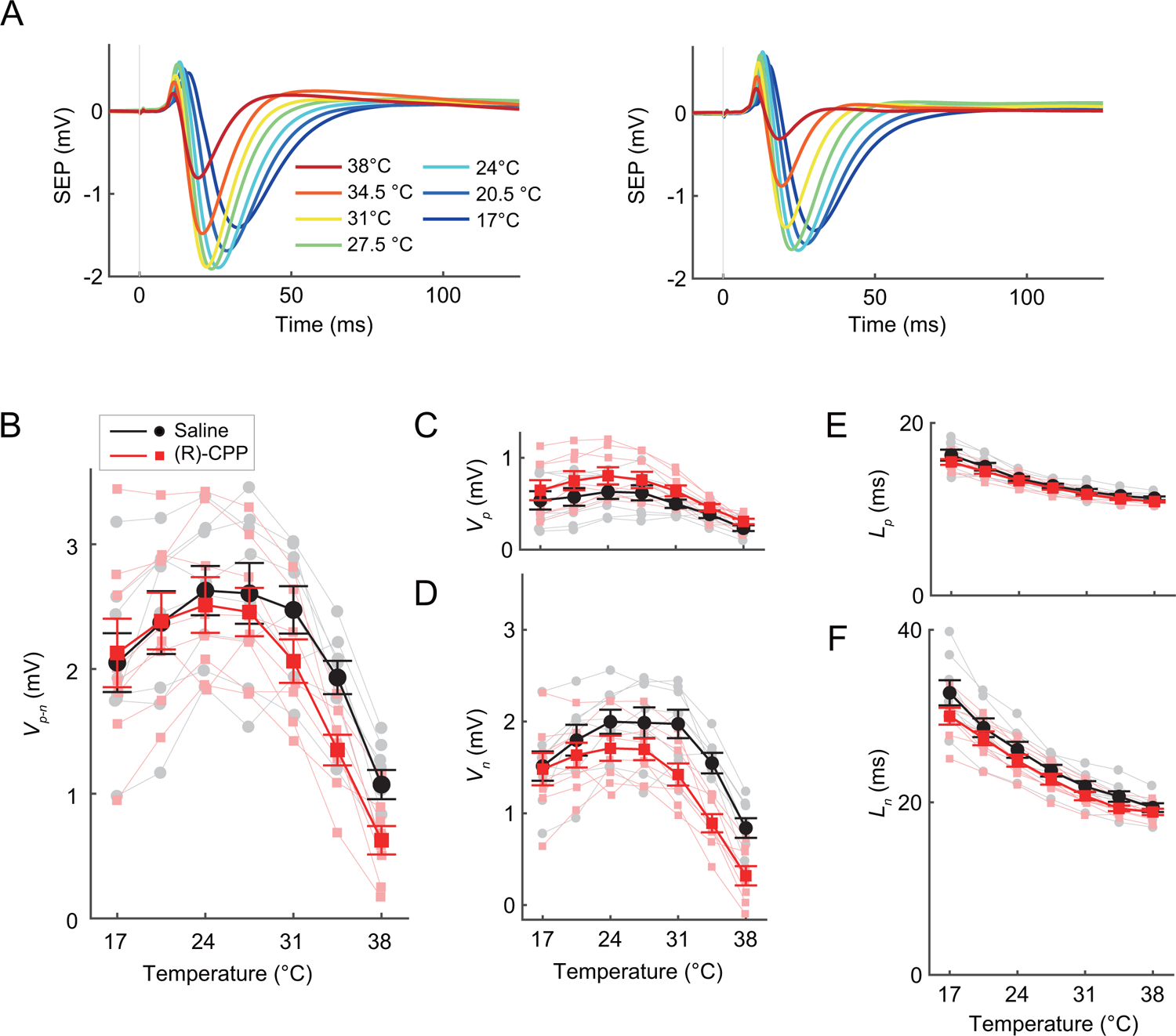
Effects of the *N*-Methyl-D-aspartic acid (NMDA) receptor antagonist (R)-CPP on the temperature dependency of somatosensory evoked potentials (SEPs) (Experiment 5). **(A)** Averaged SEP waveforms before (left panel) and after (right panel) (R)-CPP administration at various temperatures from eight animals. **(B–F)** Effects of cortical temperature on *V_p-n_* (B), *V_p_*(C), *V_n_* (D), *L_p_* (E), and *L_n_*(F) before (black circles and lines) and after (red squares and lines) (R)-CPP administration (*n* = 8 animals). Error bars indicate mean ± standard error.

**Table 9.** Results of three-way ANOVAs (antagonist by temperature and by animal) for amplitudes (*V_p-n_, V_p_*, and *V_n_*) in Experiment 5.

| Effect | Sum Sq. | *d. f.* | Mean Sq. | *F* | *p* |
| --- | --- | --- | --- | --- | --- |
| Antagonist | 1.49 | 1 | 1.49 | 33.3 | $8.5 \times 10^{-7}$ |
| Temperature | 36.27 | 6 | 6.05 | 135.2 | $7.9 \times 10^{-26}$ |
| Animal | 18.35 | 7 | 2.62 | 58.6 | $1.2 \times 10^{-19}$ |
| Antagonist×Temperature | 1.50 | 6 | 0.25 | 5.6 | $2.4 \times 10^{-4}$ |
| Antagonist×Animal | 1.53 | 7 | 0.22 | 4.9 | $4.3 \times 10^{-4}$ |
| Temperature×Animal | 9.49 | 42 | 0.23 | 5.1 | $3.3 \times 10^{-7}$ |

$$V_p$$
| Effect | Sum Sq. | *d. f.* | Mean Sq. | *F* | *p* |
| --- | --- | --- | --- | --- | --- |
| Antagonist | 0.45 | 1 | 0.45 | 120.2 | $6.7 \times 10^{-14}$ |
| Temperature | 2.50 | 6 | 0.42 | 111.4 | $3.6 \times 10^{-24}$ |
| Animal | 2.91 | 7 | 0.42 | 111.0 | $5.3 \times 10^{-25}$ |
| Antagonist×Temperature | 0.044 | 6 | 0.0073 | 2.0 | 0.094 |
| Antagonist×Animal | 0.20 | 7 | 0.029 | 7.7 | $5.7 \times 10^{-6}$ |
| Temperature×Animal | 1.35 | 42 | 0.032 | 8.6 | $8.4 \times 10^{-11}$ |

$$V_n$$
| Effect | Sum Sq. | *d. f.* | Mean Sq. | *F* | *p* |
| --- | --- | --- | --- | --- | --- |
| Antagonist | 3.57 | 1 | 3.57 | 114.6 | $1.4 \times 10^{-13}$ |
| Temperature | 20.03 | 6 | 3.34 | 107.1 | $7.9 \times 10^{-24}$ |
| Animal | 8.15 | 7 | 1.16 | 37.4 | $4.8 \times 10^{-16}$ |
| Antagonist×Temperature | 1.22 | 6 | 0.20 | 6.5 | $6.5 \times 10^{-5}$ |
| Antagonist×Animal | 0.92 | 7 | 0.13 | 4.2 | $1.3 \times 10^{-3}$ |
| Temperature×Animal | 4.35 | 42 | 0.10 | 3.3 | $8.6 \times 10^{-5}$ |

Unlike *V_p-n_*, the administration of (R)-CPP increased *V_p_* (Fig. 6C). The correlation between *V_p_* and temperature at ≥27.5 °C remained negative (*R* = −0.74, *P* < 10^−5^; Supplementary Table 16). A three-way ANOVA (antagonist by temperature and by animal) revealed significant main effects for the antagonist and temperature. The interaction between the antagonist and temperature was not significant (Table 9). Both the difference and ratio of *V_p_* before and after administering (R)-CPP (i.e., the *difference*(*V_p_*) and *ratio*(*V_p_*)) were almost constant across temperatures (Fig. S3B). The correlation between *difference*(*V_p_*) and temperature, as well as *ratio*(*V_p_*) and temperature, did not significantly deviate from 0 (*difference*(*V_p_*): *R* = −0.21, *P* = 0.12, *ratio*(*V_p_*): *R* = 0.0050, *P* = 0.97; Supplementary Tables 17–18), implying that the effect of (R)-CPP (i.e., the increase of *V_p_*) was constant across temperatures.

The administration of (R)-CPP decreased *V_n_* (Fig. 6D). The correlation between *V_n_*and temperature at ≥27.5 °C remained negative (*R* = –0.86, *P* < 10^−5^; Supplementary Table 16). A three-way ANOVA (antagonist by temperature and by animal) revealed significant main effects for the antagonist and temperature and the interaction between the antagonist and temperature (Table 9). Both the difference and ratio of *V_n_* before and after the administration of (R)-CPP (that is, the *difference*(*V_n_*) and *ratio*(*V_n_*)) were smaller at higher temperatures (Fig. S3C). The correlation between *difference*(*V_n_*) and temperature and that between *ratio*(*V_n_*) and temperature were significantly negative (*difference*(*V_n_*): *R* = −0.55, *P* = 0.000011, *ratio*(*V_n_*): *R* = −0.69, *P* < 10^−5^; Supplementary Tables 17–18), suggesting that the effect of (R)-CPP (i.e., the decrease of *V_n_*) was larger at higher temperatures.

The administration of (R)-CPP caused a slight but significant decrease in peak latencies (Fig. 6E–F). After the administration of (R)-CPP, *L_p_*maintained a negative correlation with cortical temperature but was, on average, 0.41 ms shorter. A three-way ANOVA (antagonist by temperature and by animal) revealed significant main effects for the antagonist and temperature. The interaction between the antagonist and temperature was not significant. After the administration of (R)-CPP, *L_n_*maintained a negative correlation with the cortical temperature but was, on average, 1.3 ms shorter. A three-way ANOVA (antagonist by temperature and by animal) revealed significant main effects for the antagonist and temperature, and significant interaction between the antagonist and temperature (Table 10).

**Table 10.** Results of three-way ANOVAs (antagonist by temperature and by animal) for latencies (*L_p_* and *L_n_*) in Experiment 5.

| Effect | Sum Sq. | *d. f.* | Mean Sq. | *F* | *p* |
| --- | --- | --- | --- | --- | --- |
| Antagonist | 4.66 | 1 | 4.66 | 31.8 | $1.3 \times 10^{-6}$ |
| Temperature | 301.81 | 6 | 50.30 | 342.8 | $5.2 \times 10^{-34}$ |
| Animal | 42.19 | 7 | 6.03 | 41.1 | $8.7 \times 10^{-17}$ |
| Antagonist×Temperature | 0.95 | 6 | 0.16 | 1.1 | 0.39 |
| Antagonist×Animal | 7.69 | 7 | 1.10 | 7.5 | $7.6 \times 10^{-6}$ |
| Temperature×Animal | 13.96 | 42 | 0.33 | 2.3 | $4.7 \times 10^{-3}$ |

$L_n$
| Effect | Sum Sq. | *d. f.* | Mean Sq. | *F* | *p* |
| --- | --- | --- | --- | --- | --- |
| Antagonist | 47.57 | 1 | 47.57 | 81.0 | $2.4 \times 10^{-11}$ |
| Temperature | 1920.88 | 6 | 320.15 | 545.1 | $3.6 \times 10^{-38}$ |
| Animal | 318.98 | 7 | 45.57 | 77.6 | $5.9 \times 10^{-22}$ |
| Antagonist×Temperature | 11.13 | 6 | 1.85 | 3.2 | 0.012 |
| Antagonist×Animal | 33.27 | 7 | 4.75 | 8.1 | $3.3 \times 10^{-6}$ |
| Temperature×Animal | 105.65 | 42 | 2.52 | 4.3 | $3.4 \times 10^{-6}$ |

## 3. Discussion

In the present study, we showed that the amplitudes of SEPs triggered by forepaw stimulation changed non-monotonically with the local temperature in the somatosensory cortex. For either *V_p-n_, V_n_*, or *V_p_*, the relationship between the amplitude and the temperature showed an inverted U-curve, where the SEP amplitudes increased by cooling up to approximately 27.5 °C and then decreased by further cooling (cf. Boorman et al., 2023; Gotoh et al., 2020). This temperature-dependent pattern was modulated by the administration of the GABA_A_ receptor antagonist, gabazine. The negative correlation between these amplitudes and cortical temperature at ≥27.5 °C disappeared and plateaued after gabazine administration. In addition, the negative correlation progressively disappeared as gabazine concentration increased. Notably, despite the increased SEP amplitude being associated with higher gabazine concentration, the negative correlation vanished. The effects of GABAergic inputs on SEP amplitudes, evaluated using the difference and the ratio before and after gabazine administration, were larger at higher temperatures. Note that the negative correlation at ≥27.5 °C remained after the administration of either the AMPA receptor antagonist, NBQX, or the NMDA receptor antagonist, (R)-CPP. Taken together, these results suggest that GABAergic inhibitory inputs contribute to the negative correlation between SEP amplitude and cortical temperature around the physiological cortical temperature by inducing stronger suppression of SEPs at higher temperatures.

The event-related potentials recorded from the cortical surface reflect the postsynaptic activity of pyramidal neurons, whose apical dendrites are located vertically to the cortical surface (Jackson and Bolger, 2014). Excitatory inputs to the pyramidal neurons in the deep layers create positive potentials at the surface electrode, whereas inhibitory inputs to the pyramidal neurons in the deep layers work in opposition (that is, negative potentials at the surface electrode). In contrast, excitatory inputs at the superficial layers create negative potentials, whereas inhibitory inputs at the superficial layers work oppositely (that is, positive potentials) (Jackson and Bolger, 2014). Our data showed that the administration of NBQX decreased *V_p_*and gabazine increased *V_p_*, suggesting that excitatory inputs via AMPA receptors and inhibitory inputs via GABA_A_ receptors in the deep layer contribute to the positive deflection of the SEP. In fact, the granular layer (and the infragranular layer) receive afferent inputs from the thalamus (Jackson and Bolger, 2014; Lund, 1973), and early sinks were observed in these layers in the current source density analysis of the SEPs (Jellema et al., 2004; Mitzdorf, 1985). Thus, thalamocortical inputs to the deep cortical layers may have caused the first positive deflection (P1) in our experiments.

In contrast to the AMPA receptor antagonist, the NMDA receptor antagonist, (R)-CPP, increased *V_p_*. This suggests that pyramidal neurons receive glutamatergic inputs via NMDA receptors in superficial layers during the early phase of SEPs. In fact, it has been demonstrated that the apical dendritic tufts in layer I receive input from the thalamus (Rubio-Garrido et al., 2009).

Glutamatergic inputs via NMDA receptor channels on thin dendrites generate spike-like responses called NMDA spikes (Larkum et al., 2009; Palmer, 2014), which supports the initiation of calcium spikes (Larkum et al., 2009). Thus, the present data support the possibility that glutamatergic inputs to apical dendritic tufts via NMDA receptors in layer I may contribute to the generation of the initial positive deflection (P1).

*V_n_* increased upon gabazine administration and decreased following the administration of NBQX or (R)-CPP administration. These results suggest that excitatory and inhibitory inputs to the pyramidal neurons in the superficial layers cause negative deflection (N1). The superficial layer (i.e., the supragranular layer) receives input from the granular layer within the same cortical area (Bannister, 2005; Rockland and Pandya, 1979). In addition, the supragranular layer receives feedback input from the infragranular layer of higher cortical areas (Bannister, 2005; Rockland and Pandya, 1979). These feed-forward and feedback inputs to the superficial cortical layers can generate subsequent negative deflection (N1), either directly or indirectly via excitatory and inhibitory interneurons.

The influence of the antagonists on P1/N1 deflections varied with temperature. To evaluate the contribution of excitatory and inhibitory inputs, we calculated the difference and ratio of the amplitudes before and after antagonist administration. The effects of gabazine on *V_p_, V_n_*, and *V_p-n_* (i.e., the increase in amplitude) were greater at higher temperatures. The impact of NBQX on *V_p_* (i.e., the decrease in amplitude) was smaller at higher temperatures, whereas the effects of (R)-CPP on *V_p_* (i.e., the increase in amplitude) did not show significant differences between different temperatures. The impact of NBQX on *V_n_* and *V_p-n_* did not show significant differences among the temperatures, whereas those of (R)-CPP on *V_n_* and *V_p-n_*were greater at higher temperatures than at lower temperatures. In the present study, either *V_p_, V_n_*, or *V_p-n_*showed a negative correlation with the cortical temperature at ≥27.5 °C without antagonists. Thus, the present data suggest that the larger/smaller contribution of GABAergic/glutamatergic inputs at higher temperatures leads to a negative correlation for *V_p_.* Additionally, a heightened contribution of GABAergic inputs at higher temperatures results in negative correlations for both *V_n_* and *V_p-n_*.

The temperature dependency observed in the present study was quantitatively different from that observed in the frontal cortex in a previous study (Gotoh et al., 2020). The temperature at which the amplitude was maximum was approximately 17 °C in the frontal cortex. In contrast, the temperature of the maximum amplitude was as high as 27.5 °C in the somatosensory cortex. One possible reason for this difference is the impact of inhibitory inputs that contribute to the evoked potentials. Although the proportion of rat GABAergic neurons is reported to be approximately 15% in both the frontal and somatosensory cortices, and the total number of neurons is even smaller in the frontal cortex than in the somatosensory cortex (Beaulieu, 1993), data from autoradiographic studies indicate that the densities of binding sites of [^3^H]GABA or [^3^H]muscimol to GABA_A_ receptors are higher in the frontal cortex than in the somatosensory cortex (Bowery et al., 1987; Que et al., 1999). Therefore, the contribution of GABAergic inhibitory inputs may be greater in the frontal cortex than in the somatosensory cortex. In addition, the temperature of maximum amplitude shifted toward a higher temperature than 17 °C when the concentration of gabazine increased in the frontal cortex (Gotoh et al., 2020). Therefore, we propose that the extent of the contribution of GABAergic inhibitory inputs determines the temperature-dependent pattern in the cortex and that the increased influence of inhibitory inputs lowers the temperature of the maximum amplitude in the inverted U-curve.

In the present study, focal cortical cooling also changed the SEP peak latency. Cooling the somatosensory cortex from 38–17 °C monotonically increased the P1 and N1 peak latencies. One possible reason for this delay is that cooling slows molecular processes. The slowing down of these processes can result in delays in signal conduction and synaptic transmission (Asztely et al., 1997; Cais et al., 2008; De Koninck and Mody, 1994; Esmann and Skou, 1988; Hartmann and Nothwang, 2011; Iversen and Neal, 1968; Jenkins et al., 1999; Koike and Nagata, 1979; Korogod and Demianenko, 2017; Owen et al., 2019; Postlethwaite et al., 2007; Pyott and Rosenmund, 2002; Sabatini and Regehr, 1996). In addition to delays in signal transduction, another possible reason may be the inactivation of GABAergic inhibition. A decrease in inhibition caused by focal cortical cooling may lead to a sustained excitation (Bruyns-Haylett et al., 2017; Kunori et al., 2014), resulting in a delay in the peak latency. In the present study, the application of a GABA_A_ receptor antagonist increased SEP latency, whereas the application of AMPA and NMDA receptor antagonists induced only small changes. Thus, the sustained excitation caused by disinhibition (i.e., a decrease in inhibition) at lower temperatures might have contributed to the elongated latencies in the no-antagonist conditions.

The effects of systemic changes in body temperature on SEP have been examined in clinical patients and experimental animals; such changes affect not only the local cortical temperature but also the whole somatosensory pathway from the receptors in the skin to the cortex. Thus, data describing systemic changes in body temperature include effects other than a decrease in local brain temperature, and the results have been varied. Nevertheless, data from some studies with systemic changes in body temperature might reflect the effects of the decrease in local brain temperature on SEPs. In human patients, increasing the body temperature to 42 °C (hyperthermia for the treatment of advanced neoplasms) decreases SEP amplitude (Dubois et al., 1981). In animals exposed to hyperthermia, one study showed a decrease in SEP amplitudes (< 42 °C body temperature) (Browning et al., 1992) but another study observed no significant change (38.5–39.5 °C body temperature) (Madhok et al., 2012). In these animal studies (Browning et al., 1992; Madhok et al., 2012), mild hypothermia (around 34 °C body temperature) induced an increase in SEP amplitudes, although mild hypothermia in human patients (approximately 33 °C body temperature; after cardiopulmonary resuscitation) did not elicit a consistently significant change in amplitude (Bouwes et al., 2013). Profound hypothermia (approximately 20 °C body temperature) in patients undergoing cardiothoracic surgery caused the disappearance of the cortical component peak in the SEP (N20) (Guérit et al., 1994). These studies did not contradict the inverted U relationship between amplitude and temperature reported in the present study.

In this study, we recorded SEPs by stimulating the contralateral forepaw while altering the local brain temperature. The results demonstrated that the relationship between SEP amplitude and cortical temperature was reflected by an inverted U-curve and that inhibitory inputs played a key role in determining this relationship. Recently, autism spectrum disorder (ASD), whose typical symptoms include tactile hypersensitivity, has been associated with an increase in the balance of excitation and inhibition in the cortex (Chen et al., 2020; Kondo and Lin, 2020; Lee et al., 2017; Nelson and Valakh, 2015; Rubenstein and Merzenich, 2003). Interestingly, autistic behaviors have been reported to be relieved during fever in ASD patients (Curran et al., 2007; Good, 2017), although the autistic traits evaluated using questionnaires were shown to have no significant relationship with normal body (axillary) temperatures in the general population (Hidaka et al., 2023).

It is worth noting some limitations and future directions of this study. First, the present study was performed under anesthesia. Although isoflurane is widely used in neuroscience studies, it is known not only to induce neuronal hyperpolarization but also to modulate various factors related to neurotransmitters, including GABA, glutamate, dopamine, acetylcholine, norepinephrine (He et al., 2021; Hemmings et al., 2005; Irifune et al., 1997; Leung et al., 2014; Shichino et al., 1997; Taylor et al., 2013; Uhrig et al., 2014; Zhang et al., 2020). The present data may have contaminated the influence of isoflurane anesthesia. Second, the SEPs were recorded at seven temperatures in Experiments 1, 2, 4, and 5, which could raise the question of whether some changes occurred during the long-time experiments. Although the order of cortical temperatures was counterbalanced across animals in the present study to minimize the order effects, it could still be possible that the influence of the adaptation coexisted in the present data. Third, since the antagonist administration should have been done after the control experiment, Experiments 2, 4, and 5 were always conducted after Experiment 1. This could also raise the question of whether there was an order effect of pre- and post-administration conditions. In the present study, the results showed that the effects of GABA_A_, AMPA, and NMDA receptor antagonists varied. For example, gabazine increased *V_p-n_* while NBQX and (R)-CPP decreased *V_p-n_*. Although such an order effect cannot provide an exclusive explanation for such bidirectional changes, it could still be possible that the influence of the experimental order is contaminated. Fourth, it has been widely reported that thermosensitive transient receptor potential (TRP) channels are responsive to heat and/or cold (Caterina et al., 1997; Clapham, 2003; Kashio and Tominaga, 2022; Kauer and Gibson, 2009; Zhang et al., 2023). However, the role of TRP channels in the present study has still been unknown. It is possible that these TRP channels play a role in the temperature-dependent gating in the present study. This hypothesis should be tested to understand the whole picture of the thermo-dependency of cortical activation. Fifth, the SEP amplitudes change non-lineally depending on the cortical temperature, and different antagonists affected the relationship between SEP amplitudes and temperature differently. In the present study, we did not hypothesize a specific model for fitting the SEP data in a wide range of temperatures (17−38 °C). However, it would be important to construct a mathematical model for a comprehensive understanding of the temperature dependency of neuronal activation. Finally, the body (rectal) temperature was fixed at 38 °C using a heating pad because the purpose of the study was to examine the effects of cortical temperature on SEPs. However, the brain temperature and the body temperature are interacted via heat exchange, for example, through blood flow. Thus, to understand the effects of temperature on neuronal activity, it would be important to examine whether and how the body temperature affects the brain temperature and neuronal activation. These issues should be addressed in future studies.

In conclusion, the negative correlation between SEP amplitudes and cortical temperature around the physiological cortical temperature may be caused by the high sensitivity of GABAergic inhibitory signaling to the temperature. Upon administration of the GABA_A_ receptor antagonist, gabazine, the negative correlation plateaued, and this change occurred progressively as the concentration of gabazine increased. In contrast, a negative correlation remained after administration of the AMPA receptor antagonist, NBQX, or NMDA receptor antagonist, (R)-CPP. These results suggest that GABAergic inhibitory inputs contribute to this negative correlation by suppressing SEPs to a greater extent at higher temperatures. Our findings contribute not only to the elucidation of how cortical temperature influences sensory processing under physiological conditions but also to our understanding of psychiatric disorders and establishing new treatments via future research.

## 4. Experimental Procedure

### 4.1 Animals

Thirty-two male Wistar rats (253–331 g; Japan SLC Inc., Shizuoka, Japan) were used in this study. All experimental procedures were performed in accordance with the National Institutes of Health Guide for the Care and Use of Laboratory Animals and approved by the Animal Care and Use Committee of the National Institute of Advanced Industrial Science and Technology (AIST).

### 4.2 Surgery and experimental setup

Before surgery, anesthesia was induced using 3% isoflurane (flow rate: 1.0 L/min, vehicle: room air). For electrophysiological recordings, the skull above the left somatosensory cortex was removed to create a square hole (anteroposterior, ± 2 mm; mediolateral, 2–6 mm from the bregma). The dura mater covering the exposed area was left intact. A custom-made thermal control chamber (inner diameter of approximately 5 mm) developed previously (Gotoh et al., 2020) was positioned over the skull window (Fig. 1A). The chamber can regulate cortical temperature by circulating temperature-controlled water through a stainless-steel tube. The cortical temperature and the temperature of the water circulating in the tube were linearly correlated (*R*^2^ = 0.99, *n* = 32 animals; Fig. 1B). After building the chamber, a silver ball electrode (UL-3010; Unique Medical Co., Ltd., Tokyo, Japan; diameter = 1 mm) was placed on the cortical surface 4–5 mm lateral from the midline and ± 0 mm anteroposterior to the bregma to record cortical field potentials. To monitor the cortical temperature, a thermocoupled electrode (diameter: 0.33 mm; MT-29/2; Physitemp Instruments, Inc., Clifton, NJ, USA) was inserted at a depth of 1 mm into the cortex. The thermocoupled electrode was located approximately 1.5 mm away from the ball electrode. Thirty minutes after setting up the silver ball and the thermocouple electrodes, two clip electrodes were placed on the contralateral (right) forepaw, one at the base of the fingers and the other at the wrist, to deliver electrical stimulation superficially.

### 4.3 Experiment 1: Effects of cortical temperature on SEPs

Twenty-four rats were used in Experiment 1. After filling the chamber with 100 µL saline, the cortical temperature was maintained at 38 °C for 1 h. The chamber was covered with a plastic sheet to prevent evaporation. The body (rectal) temperature was maintained at 38 °C using a heating pad (BWT-100A; BioResearch Center, Nagoya, Japan) to minimize the effects of change in the body temperature. During the experiments, anesthesia was maintained using 1.5% isoflurane (flow rate: 1.0L/min, vehicle: room air).

The cortical temperature was varied within the range of 17–38 °C in 3.5 °C steps. The SEPs were recorded in 30 trials at each temperature. To induce SEP, a single electrical pulse was delivered to the contralateral (right) forepaw (amplitude: 1 mA; duration: 1 ms). Electrical pulses were delivered every 9.9 s. The field potentials were recorded with a band-pass filter at 0.1–1,000 Hz.

The order of temperature change was counterbalanced. Twelve rats were assigned to ascending order and the other 12 rats were assigned to descending order. In the 12 rats assigned to the descending order, the initial SEP recording was conducted at 38 °C. Contrarily, in the other 12 rats assigned to the ascending order, the cortical temperature was decreased before SEP recording, and the first recording was conducted at 17 °C (approximately 30 min after the cortical temperature reached 18 °C). After each 30-trial recording, the cortical temperature was changed to the next temperature. For the cortical temperature to reach equilibrium before each recording, the recording was started ≥10 min after the temperature reached the range of ± 1 °C of the target temperature. (For example, before the experiment of 27.5 °C in the ascending [descending] order, we waited for ≥10 min after the temperature reached 26.5 °C [28.5 °C].) During this period, the cortical temperature was adjusted to reach the target temperature more precisely. After completing Experiment 1, of the 24 rats, 8 rats each participated in Experiments 2, 4, and 5, respectively.

### 4.4 Experiment 2: Effects of GABA_A_ receptor antagonist

Eight of the 24 rats used in Experiment 1 were used in Experiment 2. After Experiment 1, the saline in the chamber was replaced with 10 µM (in 100 µL saline) of the GABA_A_ receptor antagonist SR-95531 (gabazine; Toronto Research Chemicals; Toronto, ON, Canada). After incubation for 1 h at 38 °C, the SEPs were recorded using the same procedure as in Experiment 1. For each rat, the order of temperature change (i.e., ascending [*n* = 4] or descending [*n* = 4]) in Experiment 2 was consistent with that in Experiment 1.

### 4.5 Experiment 3: Effects of different concentrations of GABA_A_ receptor antagonist around the physiological temperature

Eight rats were used in Experiment 3. After filling the chamber with 100 µL saline, the cortical temperature was maintained at 38 °C for 1 h. The cortical temperature was changed within the range of 35–38 °C in 1.5 °C steps (i.e., 35, 36.5, and 38 °C). First, SEPs were recorded using a chamber filled with saline. The order of the temperature changes was counterbalanced (four for the descending order and four for the ascending order). Second, the saline in the chamber was replaced with 1 µM gabazine (in 100 µL saline). After an incubation of 1 h at 38 °C, SEPs were recorded at three temperatures. Third, the liquid in the chamber was replaced with 10 µM gabazine (in 100 µL saline). After incubation at 38 °C for 1 h, SEPs were recorded. Four rats were assigned to ascending order and the other four rats were assigned to descending order. The order of temperature change was consistent within each rat. Time intervals and other procedures followed those used in Experiments 1 and 2.

### 4.6 Experiment 4: Effects of AMPA receptor antagonist

Eight of the 24 rats used in Experiment 1 were included in Experiment 4. Following Experiment 1, the saline in the chamber was substituted with a 10 µM solution (in 100 µL saline) of the AMPA receptor antagonist NBQX disodium salt (NBQX; Tocris Bioscience; Bristol, UK). After incubation for 1 h at 38 °C, the SEPs were recorded using the same procedure as in Experiment 1. For each rat, the order of temperature change (i.e., ascending [*n* = 4] or descending [*n* = 4]) in Experiment 4 was consistent with that in Experiment 1. Time intervals and other procedures followed those used in Experiment 2.

### 4.7 Experiment 5: Effects of NMDA receptor antagonist

Eight of the 24 rats used in Experiment 1 were included in Experiment 5. Following Experiment 1, the saline in the chamber was substituted with a 10 µM solution (in 100 µL saline) of the NMDA receptor antagonist (R)-CPP (Tocris Bioscience; Bristol, UK). After incubation for 1 h at 38 °C, the SEPs were recorded using the same procedure as in Experiment 1. For each rat, the order of temperature change (i.e., ascending [*n* = 4] or descending [*n* = 4]) in Experiment 5 was consistent with that in Experiment 1. Time intervals and other procedures followed those used in Experiment 2.

### 4.8 Data analysis

The SEP data between −1,000 and 200 ms from the onset of the electrical pulse (0 ms) were used for the data analysis. The SEP data for each trial were calibrated by subtracting an average of the SEP data between −50 and 0 ms to obtain the baseline (0 mV). According to a preliminary analysis, SEPs were sometimes disturbed by spontaneous activity close to or during the evoked response. To precisely evaluate the effect of temperature on SEPs, data from trials in which spontaneous activity was not observed close to or during stimulation were analyzed.

First, the data of trials in which the difference between maximum and minimum values in any 25-ms time bin between −1,000 and 0 ms exceeded 1.0 mV were excluded from the analyses (1,022 of 12,240 trials). Second, the data of trials in which the absolute values between 2 and 7 ms (i.e., for 5 ms after the electrical noise disappeared) exceeded 0.5 mV were excluded from the analyses (36 of the remaining 11,218 trials). In total, data from 1,058 of 12,240 (8.6%) trials were excluded from the analyses. Following exclusion, the SEP data for each temperature in each animal were averaged.

SEP indices were calculated for the averaged SEP data at each temperature for each animal (Fig. 2B). The averaged data were smoothed using a moving-average filter with five-time bins (approximately 0.4 ms). To automatically find the peaks of P1 and N1 deflections, we used the following algorithm: 1) Find the local maximums during 2–20 ms (2–30 ms in Experiment 2) after stimulation. 2) The largest local maximum was defined as the P1 peak. 3) Find the first local minimum ≥2 ms after the P1 peak. 4) Find the local minimums during 0–5 ms after the first local minimum. 5) The smallest local minimum was defined as the N1 peak. The P1 peak amplitude (*V_p_*) was defined as the voltage at P1 and the N1 peak amplitude (*V_n_*) was defined as the sign-inverted voltage at N1. Peak-to-peak amplitude (*V_p-n_*) was defined as the difference between P1 and N1. The P1 (*L_p_*) and N1 peak latencies (*L_n_*) were defined as the time between the onset of stimulation (0 ms) and the respective peaks.

To examine temperature dependency on *V_p_, V_n_*, and *V_p-n_*, we calculated correlation coefficients between these amplitudes and cortical temperatures in the range of ≤27.5 °C and ≥27.5 °C, separately, in Experiments 1, 2, 4, and 5. To examine the effects of cortical temperature on the amplitudes and latencies with no antagonist, we performed a two-way ANOVA (temperature by animal) for the amplitudes and latencies in Experiment 1. To examine the effects of cortical temperature and antagonists, we performed three-way ANOVAs (antagonist by temperature and by animal) for the amplitudes and latencies in Experiment 2–5. To examine the effects of antagonists on the amplitudes at each temperature, we calculated the difference and ratio between the amplitudes before (Experiment 1) and after antagonist application (Experiments 2, 4, and 5) as follows:

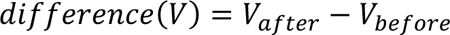

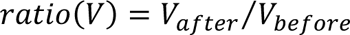

where *V* denotes the amplitude (*V_p_, V_n_*, or *V_p-n_*), and *V_before_* and *V_after_* denote the amplitudes before and after the antagonist application, respectively. For these differences and ratios, we calculated the correlation coefficients with the temperature.

In Experiment 3, the amplitudes and latencies were fitted using the following linear function for each rat:

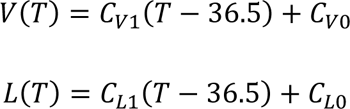

where *T* denotes the cortical temperature, *V* and *L* denote the amplitude and the latency, *C_V1_* and *C_L1_*denote the slope of the amplitude and the latency, and *C_V0_* and *C_L0_* denote the constant term (that is, the estimated amplitudes and latencies at 36.5 °C), respectively. We also calculated the change ratios *CR_V_* and *CR_L_*(i.e., the percentage of the slope to the estimated amplitude and latency at 36.5 °C, respectively):

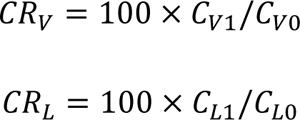

Tests for the null hypothesis that the slope is equal to zero and that the change ratio is equal to zero were performed using a *t*-test. Additionally, we performed two-way ANOVAs (antagonist by animal) for *C_V0_, C_V1_, C_L0_, C_L1_, CR_V_*, and *CR_L_*. MATLAB (R2020b, The MathWorks, Inc., Natick, MA, USA) was used for all data analyses.

## Acknowledgments

We thank Shigeru Yamane, Yoshihisa Tachibana, and Makoto Wada for their valuable discussions.

## Footnotes

1 FBC, focal brain cooling; SEPs, somatosensory evoked potentials; P1, positive deflection; N1, negative deflection; AMPA, aminomethylphosphonic acid; NMDA, *N*-Methyl-D-aspartic acid; GABA, gamma-aminobutyric acid; GABA_A_, gamma-aminobutyric acid type A

