## Supplementary Materials for "Effects of focal cortical cooling on somatosensory evoked potentials in rats"

### Figure S1

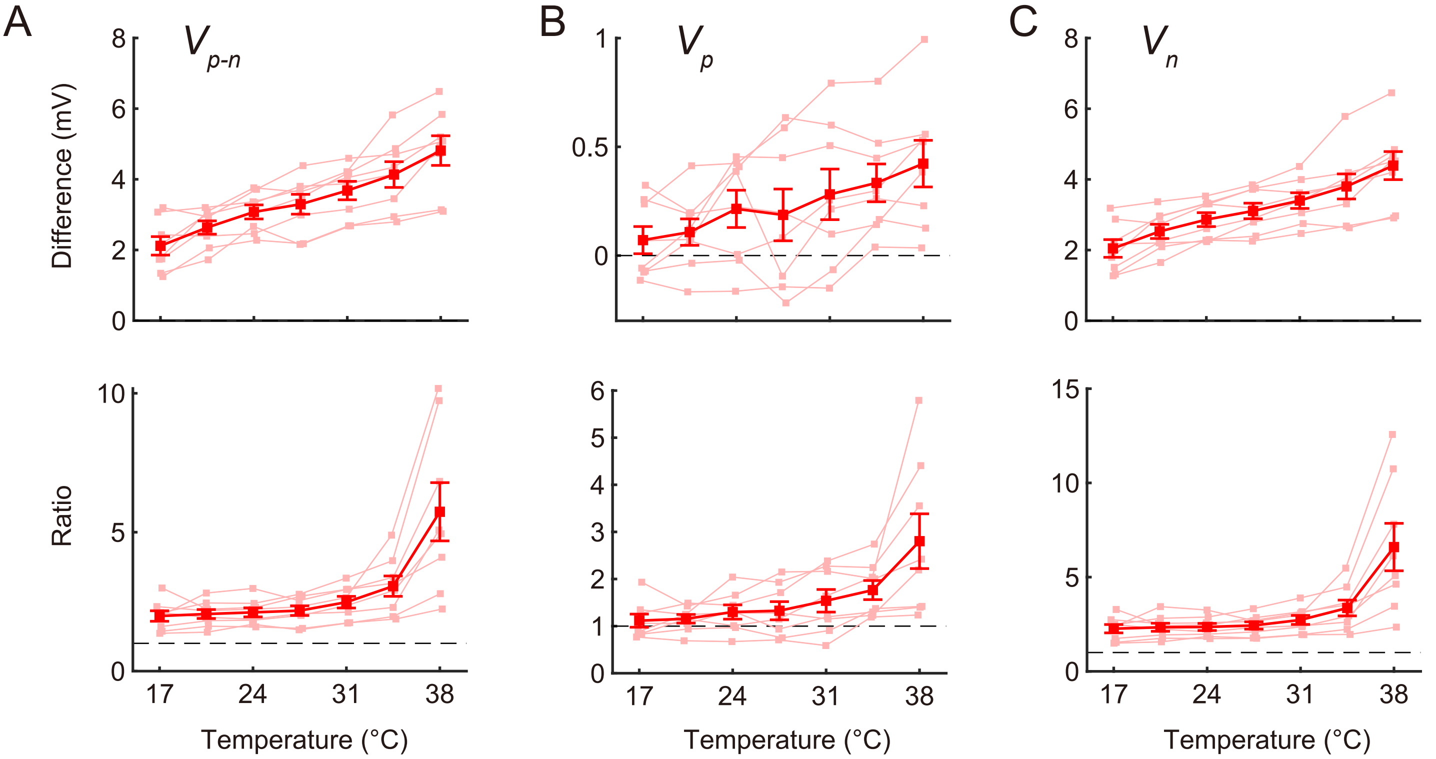


**Figure S1.** Comparison of amplitudes before and after gamma-aminobutyric acid type A ($GABA_A$) receptor antagonist gabazine administration. Differences (top panels) and ratios (bottom panels) of amplitudes (A: $V_{p\text{-}n}$, B: $V_p$, C: $V_n$) before and after gabazine administration were plotted against cortical temperature (n = 8 animals). Error bars indicate mean ± standard error.

### Figure S2

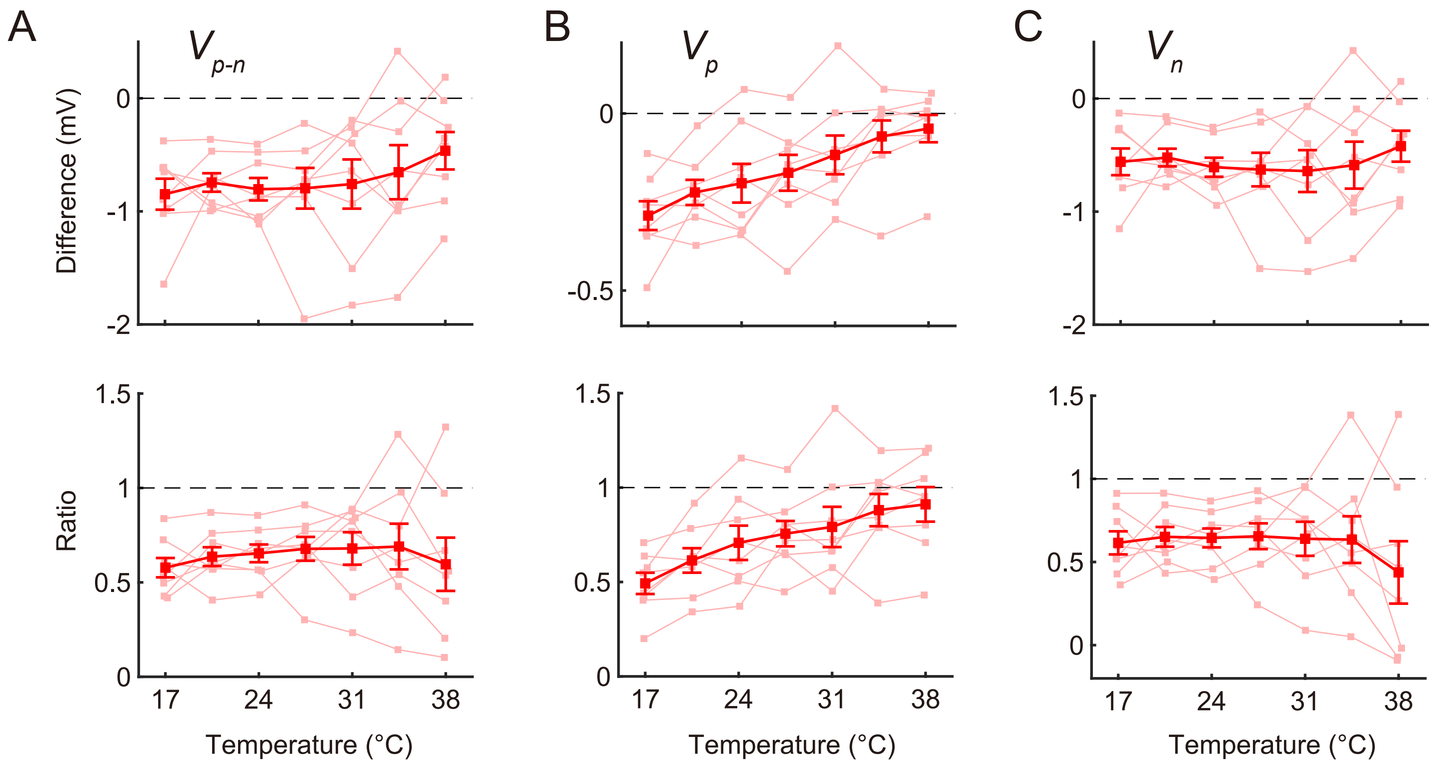


**Figure S2.** Comparison of amplitudes before and after aminomethylphosphonic acid (AMPA) receptor antagonist NBQX administration. Differences (top panels) and ratios (bottom panels) of amplitudes (A: $V_{p-n}$, B: $V_p$, C: $V_n$) before and after NBQX administration were plotted against cortical temperature (n = 8 animals). Error bars indicate mean ± standard error.

### Figure S3

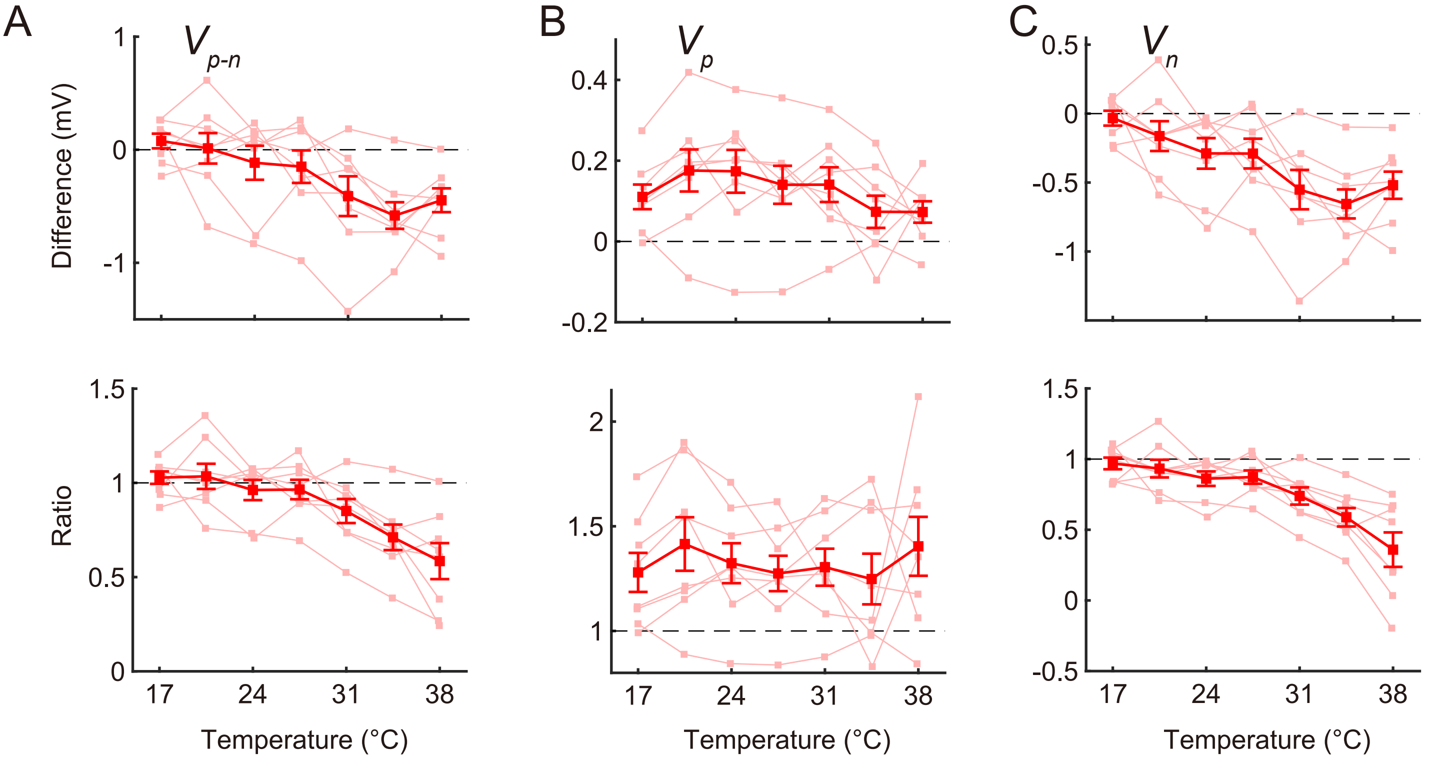


**Figure S3.** Comparison of amplitudes before and after *N*-Methyl-D-aspartic acid (NMDA) receptor antagonist (R)-CPP administration. Differences (top panels) and ratios (bottom panels) of amplitudes (A: $V_{p\text{-}n}$, B: $V_p$, C: $V_n$) before and after (R)-CPP administration were plotted against cortical temperature (n = 8 animals). Error bars indicate mean ± standard error.

**Supplementary Table 1.** Results of correlation analyses between temperatures (≤27.5 °C and ≥27.5°C) and amplitudes ($V_{p\text{-}n}$, $V_p$, and $V_n$) in Experiment 1.

| | ≦27.5 °C | | ≧27.5 °C | |
|---|---|---|---|---|
| | *r* | *p* | *r* | *p* |
| $V_{p\text{-}n}$ | 0.31 | $1.9\times10^{-3}$ | −0.70 | $1.9\times10^{-15}$ |
| $V_p$ | 0.15 | 0.15 | −0.65 | $9.6\times10^{-13}$ |
| $V_p$ | 0.36 | $3.4\times10^{-4}$ | −0.67 | $5.9\times10^{-14}$ |

**Supplementary Table 2.** Results of correlation analyses between temperatures (≤27.5 °C and ≥27.5°C) and amplitudes ($V_{p\text{-}n}$, $V_p$, and $V_n$) before and after gabazine administration in Experiment 2.

| | | ≦27.5 °C | | ≧27.5 °C | |
|---|---|---|---|---|---|
| | | *r* | *p* | *r* | *p* |
| $V_{p\text{-}n}$ | No antagonist | 0.33 | 0.067 | −0.66 | $3.4\times10^{-5}$ |
| | Gabazine | 0.81 | $1.5\times10^{-8}$ | −0.11 | 0.54 |
| $V_p$ | No antagonist | 0.17 | 0.36 | −0.58 | $4.5\times10^{-4}$ |
| | Gabazine | 0.26 | 0.15 | −0.23 | 0.21 |
| $V_n$ | No antagonist | 0.37 | 0.037 | −0.65 | $6.3\times10^{-5}$ |
| | Gabazine | 0.79 | $8.1\times10^{-8}$ | 0.010 | 0.96 |

**Supplementary Table 3.** Results of correlation analyses between temperature and the difference between amplitudes ($V_{p\text{-}n}$, $V_p$, and $V_n$) before and after gabazine administration in Experiment 2.

| | $r$ | $p$ |
|---|---|---|
| *difference*($V_{p\text{-}n}$) | 0.73 | $1.7\times10^{-10}$ |
| *difference*($V_p$) | 0.41 | $1.7\times10^{-3}$ |
| *difference*($V_n$) | 0.70 | $1.4\times10^{-9}$ |

**Supplementary Table 4.** Results of correlation analyses between temperature and the ratio of amplitudes ($V_{p\text{-}n}$, $V_p$, and $V_n$) before and after gabazine administration in Experiment 2.

| | $r$ | $p$ |
|---|---|---|
| $ratio(V_{p\text{-}n})$ | 0.57 | $5.2\times10^{-6}$ |
| $ratio(V_p)$ | 0.51 | $5.0\times10^{-5}$ |
| $ratio(V_n)$ | 0.55 | $1.4\times10^{-5}$ |

**Supplementary Table 5.** Results of the linear model fitting for $V_{p\text{-}n}$, $V_p$, and $V_n$ in Experiment 3.

| | | Constant term | | Slope | | Change ratio | |
|---|---|---|---|---|---|---|---|
| | | $C_0$ | $p$ | $C_1$ | $p$ | $CR_V$ | $p$ |
| $V_{p\text{-}n}$ | 0 μM | 1.17 | $1.6\times10^{-6}$ | −0.28 | $6.6\times10^{-4}$ | −25.02 | $1.2\times10^{-3}$ |
| | 1 μM | 3.48 | $3.4\times10^{-5}$ | −0.20 | 0.015 | −7.34 | 0.021 |
| | 10 μM | 6.08 | $5.7\times10^{-8}$ | −0.0065 | 0.83 | −0.16 | 0.75 |
| $V_p$ | 0 μM | 0.19 | $1.6\times10^{-3}$ | −0.042 | $2.0\times10^{-3}$ | −35.23 | 0.023 |
| | 1 μM | 0.47 | $2.6\times10^{-5}$ | −0.046 | $7.7\times10^{-3}$ | −10.34 | 0.015 |
| | 10 μM | 0.69 | $5.9\times10^{-5}$ | −0.0084 | 0.39 | −0.63 | 0.72 |
| $V_n$ | 0 μM | 0.97 | $1.9\times10^{-6}$ | −0.24 | $6.9\times10^{-4}$ | −25.64 | $8.3\times10^{-4}$ |
| | 1 μM | 3.01 | $7.1\times10^{-5}$ | −0.16 | 0.025 | −6.90 | 0.027 |
| | 10 μM | 5.39 | $1.7\times10^{-7}$ | 0.0019 | 0.94 | −0.022 | 0.96 |

**Supplementary Table 6.** Results of two-way ANOVAs (antagonist by animal) for the constant terms (i.e., the estimated amplitudes at 36.5 °C) of the linear models for $V_{p\text{-}n}$, $V_p$, and $V_n$ in Experiment 3.

$V_{p\text{-}n}$

| Effect | Sum Sq. | *d.f.* | Mean Sq. | *F* | *p* |
|---|---|---|---|---|---|
| Antagonist | 96.61 | 2 | 48.30 | 99.3 | $5.4\times10^{-9}$ |
| Animal | 4.98 | 7 | 0.71 | 1.5 | 0.26 |

$V_p$

| Effect | Sum Sq. | *d.f.* | Mean Sq. | *F* | *p* |
|---|---|---|---|---|---|
| Antagonist | 0.99 | 2 | 0.50 | 43.8 | $9.5\times10^{-7}$ |
| Animal | 0.42 | 7 | 0.060 | 5.3 | $3.9\times10^{-3}$ |

$V_n$

| Effect | Sum Sq. | *d.f.* | Mean Sq. | *F* | *p* |
|---|---|---|---|---|---|
| Antagonist | 78.10 | 2 | 39.05 | 95.9 | $6.8\times10^{-9}$ |
| Animal | 5.85 | 7 | 0.84 | 2.1 | 0.12 |

**Supplementary Table 7.** Results of two-way ANOVAs (antagonist by animal) for the slope of the linear models for $V_{p\text{-}n}$, $V_p$, and $V_n$ in Experiment 3.

$V_{p\text{-}n}$

| Effect | Sum Sq. | *d.f.* | Mean Sq. | *F* | *p* |
|---|---|---|---|---|---|
| Antagonist | 0.32 | 2 | 0.16 | 10.4 | $1.7\times10^{-3}$ |
| Animal | 0.19 | 7 | 0.028 | 1.8 | 0.16 |

$V_p$

| Effect | Sum Sq. | *d.f.* | Mean Sq. | *F* | *p* |
|---|---|---|---|---|---|
| Antagonist | 0.0067 | 2 | 0.0033 | 4.9 | 0.024 |
| Animal | 0.0081 | 7 | 0.0012 | 1.7 | 0.19 |

$V_n$

| Effect | Sum Sq. | *d.f.* | Mean Sq. | *F* | *p* |
|---|---|---|---|---|---|
| Antagonist | 0.24 | 2 | 0.12 | 10.3 | $1.8\times10^{-3}$ |
| Animal | 0.14 | 7 | 0.019 | 1.7 | 0.20 |

**Supplementary Table 8.** Results of two-way ANOVAs (antagonist by animal) for the change ratio of the linear models for $V_{p\text{-}n}$, $V_p$, and $V_n$ in Experiment 3.

$V_{p\text{-}n}$

| Effect | Sum Sq. | *d. f.* | Mean Sq. | *F* | *p* |
|---|---|---|---|---|---|
| Antagonist | 2620.71 | 2 | 1310.36 | 18.4 | $1.2\times10^{-4}$ |
| Animal | 625.30 | 7 | 89.33 | 1.3 | 0.34 |

$V_p$

| Effect | Sum Sq. | *d. f.* | Mean Sq. | *F* | *p* |
|---|---|---|---|---|---|
| Antagonist | 5096.76 | 2 | 2548.38 | 5.2 | 0.021 |
| Animal | 2041.95 | 7 | 291.71 | 0.6 | 0.75 |

$V_n$

| Effect | Sum Sq. | *d. f.* | Mean Sq. | *F* | *p* |
|---|---|---|---|---|---|
| Antagonist | 2812.37 | 2 | 1406.19 | 22.0 | $4.8\times10^{-5}$ |
| Animal | 634.98 | 7 | 90.71 | 1.4 | 0.27 |

**Supplementary Table 9.** Results of the linear model fitting for $L_p$ and $L_n$ in Experiment 3.

| | | Constant term | | Slope | | Change ratio | |
|---|---|---|---|---|---|---|---|
| | | $C_0$ | $p$ | $C_1$ | $p$ | $CR_L$ | $p$ |
| $L_p$ | 0 μM | 10.36 | $2.1\times10^{-8}$ | −0.28 | 0.15 | −3.21 | 0.15 |
| | 1 μM | 10.84 | $1.0\times10^{-12}$ | −0.092 | 0.18 | −0.84 | 0.18 |
| | 10 μM | 12.82 | $2.1\times10^{-8}$ | −0.33 | $1.4\times10^{-4}$ | −2.58 | $6.6\times10^{-5}$ |
| $L_n$ | 0 μM | 18.56 | $7.8\times10^{-11}$ | −0.28 | $5.3\times10^{-3}$ | −1.49 | $5.0\times10^{-3}$ |
| | 1 μM | 20.50 | $8.3\times10^{-9}$ | −0.34 | 0.019 | −1.77 | 0.020 |
| | 10 μM | 28.98 | $1.3\times10^{-7}$ | −0.94 | $7.4\times10^{-4}$ | −3.17 | $1.5\times10^{-4}$ |

**Supplementary Table 10.** Results of two-way ANOVAs (antagonist by animal) for the constant terms (i.e., the estimated latencies at 36.5 °C) of the linear models for $L_p$ and $L_n$ in Experiment 3.

$L_p$

| Effect | Sum Sq. | *d.f.* | Mean Sq. | *F* | *p* |
|---|---|---|---|---|---|
| Antagonist | 27.21 | 2 | 13.60 | 10.2 | $1.9\times10^{-3}$ |
| Animal | 1.88 | 7 | 0.27 | 0.2 | 0.98 |

$L_n$

| Effect | Sum Sq. | *d.f.* | Mean Sq. | *F* | *p* |
|---|---|---|---|---|---|
| Antagonist | 491.51 | 2 | 245.76 | 57.5 | $1.8\times10^{-7}$ |
| Animal | 73.19 | 7 | 10.46 | 2.4 | 0.073 |

**Supplementary Table 11.** Results of two-way ANOVAs (antagonist by animal) for the slope of the linear models for $L_p$ and $L_n$ in Experiment 3.

$L_p$

| Effect | Sum Sq. | *d. f.* | Mean Sq. | *F* | *p* |
|---|---|---|---|---|---|
| Antagonist | 0.26 | 2 | 0.13 | 1.4 | 0.29 |
| Animal | 0.67 | 7 | 0.10 | 1.0 | 0.46 |

$L_n$

| Effect | Sum Sq. | *d. f.* | Mean Sq. | *F* | *p* |
|---|---|---|---|---|---|
| Antagonist | 2.11 | 2 | 1.06 | 7.5 | $6.1\times10^{-3}$ |
| Animal | 0.53 | 7 | 0.08 | 0.5 | 0.79 |

**Supplementary Table 12.** Results of two-way ANOVAs (antagonist by animal) for the change ratio of the linear models for $L_p$ and $L_n$ in Experiment 3.

$L_p$

| Effect | Sum Sq. | *d. f.* | Mean Sq. | *F* | *p* |
|---|---|---|---|---|---|
| Antagonist | 23.96 | 2 | 11.98 | 1.0 | 0.40 |
| Animal | 75.22 | 7 | 10.75 | 0.9 | 0.56 |

$L_n$

| Effect | Sum Sq. | *d. f.* | Mean Sq. | *F* | *p* |
|---|---|---|---|---|---|
| Antagonist | 13.07 | 2 | 6.53 | 2.9 | 0.085 |
| Animal | 6.72 | 7 | 0.96 | 0.4 | 0.87 |

**Supplementary Table 13.** Results of correlation analyses between temperatures (≤27.5 °C and ≥27.5 °C) and amplitudes ($V_{p\text{-}n}$, $V_p$, and $V_n$) before and after NBQX administration in Experiment 4.

| | | ≦27.5 °C | | ≧27.5 °C | |
|---|---|---|---|---|---|
| | | *r* | *p* | *r* | *p* |
| $V_{p\text{-}n}$ | No antagonist | 0.36 | 0.041 | −0.80 | $3.5\times10^{-8}$ |
| | NBQX | 0.34 | 0.053 | −0.69 | $1.3\times10^{-5}$ |
| $V_p$ | No antagonist | 0.13 | 0.47 | −0.77 | $2.1\times10^{-7}$ |
| | NBQX | 0.42 | 0.017 | −0.63 | $9.5\times10^{-5}$ |
| $V_n$ | No antagonist | 0.42 | 0.015 | −0.76 | $5.9\times10^{-7}$ |
| | NBQX | 0.28 | 0.12 | −0.63 | $9.7\times10^{-5}$ |

**Supplementary Table 14.** Results of correlation analyses between temperature and the differences between amplitudes ($V_{p\text{-}n}$, $V_p$, and $V_n$) before and after NBQX administration in Experiment 4.

| | $r$ | $p$ |
|---|---|---|
| $difference(V_{p\text{-}n})$ | 0.21 | 0.12 |
| $difference(V_p)$ | 0.55 | $1.1\times10^{-5}$ |
| $difference(V_n)$ | 0.045 | 0.74 |

**Supplementary Table 15.** Results of correlation analyses between temperature and the ratio between amplitudes ($V_{p\text{-}n}$, $V_p$, and $V_n$) before and after NBQX administration in Experiment 4.

| | $r$ | $p$ |
|---|---|---|
| $ratio(V_{p\text{-}n})$ | 0.057 | 0.68 |
| $ratio(V_p)$ | 0.52 | $3.8\times10^{-5}$ |
| $ratio(V_n)$ | −0.14 | 0.31 |

**Supplementary Table 16.** Results of correlation analyses between temperatures (≤27.5 °C and ≥27.5 °C) and amplitudes ($V_{p\text{-}n}$, $V_p$, and $V_n$) before and after (R)-CPP administration in Experiment 5.

| | | ≦27.5 °C | | ≧27.5 °C | |
|---|---|---|---|---|---|
| | | $r$ | $p$ | $r$ | $p$ |
| $V_{p\text{-}n}$ | No antagonist | 0.33 | $6.6\times10^{-2}$ | −0.75 | $8.5\times10^{-7}$ |
| | (R)-CPP | 0.20 | 0.28 | −0.86 | $4.6\times10^{-10}$ |
| $V_p$ | No antagonist | 0.15 | 0.43 | −0.70 | $7.4\times10^{-6}$ |
| | (R)-CPP | 0.16 | 0.37 | −0.74 | $1.2\times10^{-6}$ |
| $V_n$ | No antagonist | 0.40 | 0.024 | −0.73 | $2.3\times10^{-6}$ |
| | (R)-CPP | 0.21 | 0.26 | −0.86 | $1.9\times10^{-10}$ |

**Supplementary Table 17.** Results of correlation analyses between temperature and the differences between amplitudes ($V_{p\text{-}n}$, $V_p$, and $V_n$) before and after (R)-CPP administration in Experiment 5.

| | $r$ | $p$ |
|---|---|---|
| *difference*($V_{p\text{-}n}$) | −0.52 | $3.8\times10^{-5}$ |
| *difference*($V_p$) | −0.21 | 0.12 |
| *difference*($V_n$) | −0.55 | $1.1\times10^{-5}$ |

**Supplementary Table 18.** Results of correlation analyses between temperature and the ratio of amplitudes ($V_{p\text{-}n}$, $V_p$, and $V_n$) before and after (R)-CPP administration in Experiment 5.

| | $r$ | $p$ |
|---|---|---|
| $ratio(V_{p\text{-}n})$ | −0.64 | $1.0\times10^{-7}$ |
| $ratio(V_p)$ | 0.0050 | 0.97 |
| $ratio(V_n)$ | −0.69 | $3.5\times10^{-9}$ |
